# Self-consistent migration puts tight constraints on the spatio-temporal organization of species-rich metacommunities

**DOI:** 10.1101/2021.12.14.472702

**Authors:** Jonas Denk, Oskar Hallatschek

## Abstract

Biodiversity is often attributed to a dynamic equilibrium between immigration of species and competition-driven extinction. This equilibrium forms a common basis for studying ecosystem assembly from a static reservoir of migrants–the mainland. Yet, natural ecosystems often consist of many coupled communities (i.e. metacommunities) and migration occurs *between* these communities. The pool of migrants then depends on what is sustained in the ecosystem, which in turn depends on the dynamic migrant pool. This chicken-and-egg problem of survival and migration is poorly understood in communities of many competing species, except for the neutral case - the “unified neutral theory of biodiversity”. Employing spatio-temporal simulations and mean-field analyses, we show that self-consistent migration puts rather tight constraints on the dynamic migration-extinction equilibrium. When the number of species is large, even weak competitions push species to the edge of their global extinction, such that the overall diversity is highly sensitive to perturbations in demographic parameters, including growth and dispersal rates. When migration is short-range, the resulting spatiotemporal abundance patterns follow broad scale-free distributions that correspond to a directed percolation phase transition. The qualitative agreement of our results for short-range and long-range migration suggests that this self-organization process is a general property of species-rich metacommunities. Our study shows that self-sustaining metacommunities are highly sensitive to environmental change and provides insights into how biodiversity can be rescued and maintained.

---

The dynamics of an ecological community is shaped by the interplay of numerous factors, including inter- and intraspecies interactions, speciation and species immigration. Finding meaningful theoretical models for assembly and stability of ecosystems is further complicated by the overwhelming number of species typically found in natural ecosystems (1–6). Despite this complexity, insights into some statistical properties of the ecosystem can be gained by assuming a dynamic equilibrium between extinction of species in a local community (island) and immigration of species from some static reservoir (mainland)–a concept that builds on MacArthur and Wilson’s classical theory of island biography (7). For instance, neutral theories (8), where differences in the individuals’ interactions are neglected and extinctions occur due to demographic stochasticity, have shown that the balance between extinctions and continuous immigration leads to surprisingly good estimates of naturally ob-served quantities including static species abundance distributions (9, 10). In an attempt to account for differences in species interactions, theoretical studies (11–17) have considered randomly-distributed interactions among species and have found that when the number of species on an island is large, some of the macroscopic properties of the ecosystem, such as its stability and abundance distribution, can be understood based on the interaction statistics alone.

While in mainland-island models the dynamic equilibrium of a local community strongly depends on migrants from a static mainland, it is natural to ask how biodiversity can be maintained when migrants instead come from other local communities themselves. Natural ecosystems, for instance, are often better represented as *metacommunities* composed of many coupled communities, between which individuals migrate (18, 19). The pool of migrants then depends on what is sustained in the ecosystem, which in turn depends on the migrant pool.

Spatially extended ecosystem - prime examples of such meta-communities - can vary over widely different length scales ranging from microbial biofilms to tropical forests (18). In biofilms, for instance, the interplay of dispersal and interspecies competition can lead to intriguing spatial structure with important impacts on the stability and evolution of the metacommunity (20–24). Over the past 50 years, theoretical studies on metacommunities (25–34) have repeatedly shown that self-consistent migration between patches can alleviate global extinctions of species. Simply put, migration can prevent global extinction because even if species go extinct on some patches, they can still be present on other patches and from there recolonize the patches where they had gone extinct. When inter-species interactions are strong and cause large fluctuations in the species’ abundances (e.g. through predator-prey interactions), migration within a meta-community can lead to intriguing spatio-temporal abundance patterns. These patterns may include traveling waves (31) and–when the number of species is large–chaotic dynamics (33, 34). The alternative regime where inter-species competitions are weak (relative to intra-species competitions) occurs when species occupy very different niches (35, 36), as proposed for various natural ecosystems (4, 37–41). In this case, intrinsic fluctuations due to the species’ interactions are strongly suppressed (17, 35, 36) and become overshadowed by demographic fluctuations as more and more species are packed into the ecosystem.

Here, we develop stochastic approaches to explore how self-consistent migration shapes such species-rich communities. We find that, as species numbers increase, increasing local demographic fluctuations occurring within species drive the system to a migration-dependent edge of global extinction, leading to properties that are independent of the microscopic details of our model. Motivated by the large variation of migration length scales in natural ecosystems [e.g. compare the movement of microbes in biofilms and long-distance dispersal of seeds (42)], we consider migration on two limiting length-scales: short-range migration between nearest neighboring patches and spatially uniform migration between all patches (global migration). For short-range migration, we find that the proximity of species to their critical threshold results in fractal spatio-temporal patterns, reminiscent of patterns close to a directed percolation threshold. For global migration, we derive an analytical mean-field approximation for the abundance distribution. Based on this meanfield approach, we can relate equilibrium properties of our model to earlier ecological theories without metacommunity structure. Our study sheds new light on spatially structured metacommunities and suggests that self-consistent migration renders species-rich metacommunity much more sensitive to perturbations, including environmental change, than previously thought.

## Results

### Lotka-Volterra model of metacommunities with weak inter-species competition

In the following, we consider *S* species that live in a metacommunity of *P* coupled communities (patches), where *P* is assumed to be large. The dynamics of the species’ populations is modeled by the following set of generalized Lotka-Volterra equations (see Fig. 1 for a graphical representation):

**Fig. 1.**
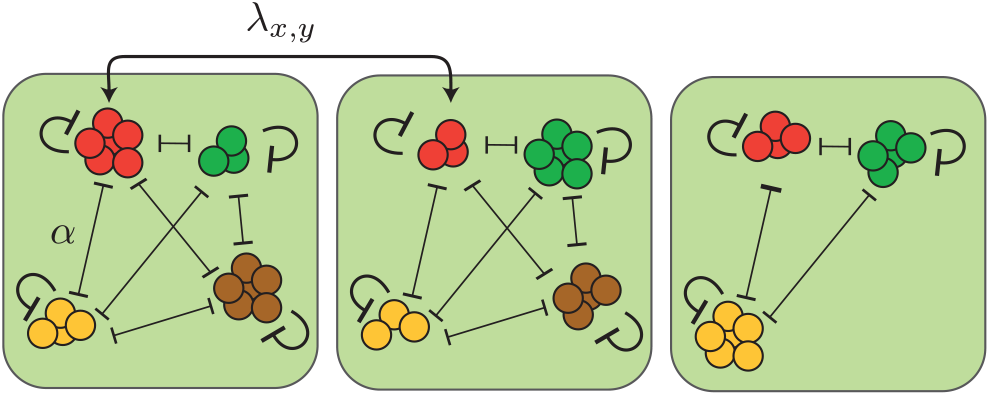
Metacommunity framework with niche interactions. Populations of different species (circles illustrate individuals) grow on patches (green) with growth rate *r* and carrying capacity *K*. Competition between species, characterized by the competition strength *α* are weaker than self-limitation, e.g. due to limited niche overlap, allowing multiple species to coexist on a patch. Furthermore, individuals migrate between different patches *x* and *y* at migration rate *λ*_*x,y*_. Especially when a species’ population size on a patch is low (e.g. due to a large number of competitors), demographic fluctuations promote stochastic extinctions of species on individual patches (see extinction of the brown species on the right patch).

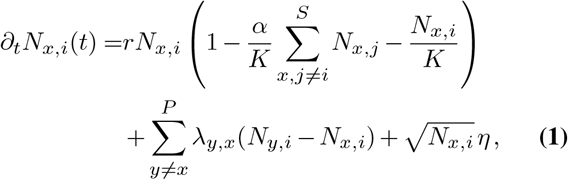

where *N*_*x,i*_ denotes the abundance of species *i* ∈{1, …*S*} on the patch *x* ∈{1, …, *P*}. The first term in Eq. (1) describes growth of a species’ population at a growth rate *r*, which is bounded by self-limiting interactions within a species as well as competition with all other species on the same patch. For a clearer presentation of our main results, the strengths of interspecies interactions is chosen to be identical for all species and set to *α*. Later, we relax this assumption and will allow variations in the species’ inter-species interactions, growth rates and migration rates. In the absence of inter-species interactions (i.e. *α* = 0), self-limiting interactions lead to population saturation at a carrying capacity *K*. Thus, *α* can be interpreted as the ratio of inter-species and self-limiting interaction strengths. By setting 0 *< α <* 1 we assume that self-limiting interactions are stronger than competition between species. This assumption emulates ecosystems where species coexist by occupying different niches, as frequently suggested for natural microbial ecosystems (4, 40) [*α* could be interpreted as a measure for the niche overlap (43, 44)]. In the following we focus on weak competition and choose 0 *< α*≪1, which will allow multiple species to coexist on each patch. On the other hand, strong inter-species competition (*α >* 1) is known to promote exclusion between species (43, 45). The special case *α* = 1 marks the neutral scenario (8), and sets the boundary between niche partitioning (0 *< α <* 1) and competitive exclusion (*α >* 1). The second term in Eq. (1) takes into account migration, where *λ*_*x,y*_ denotes the migration rate between two patches *x* and *y* and is assumed equal for all species (we will later relax the assumption of equal migration rates). The last term in Eq. (1) reflects demographic fluctuations due to random births and deaths of individuals within a population (46). Here, *η*_*x,i*_ denotes uncorrelated noise with zero mean and variance *ω*^2^. Assuming 0 *< α <* 1, the deterministic dynamics of Eq. (1) (i.e. ignoring noise) possess a stable solution in which all species coexist on all patches at equal abundance *N* ^*^ with

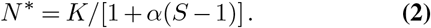

The inverse dependence of *N* ^*^ on the number of species *S* suggests that when the number of species is large, the population size of each species on a patch can become very small, favoring (local) stochastic extinctions of species (compare extinction of brown species illustrated in Fig. 1). This leads us to suspect that, especially in the case of many coexisting species, migration plays an important role in offsetting local species extinctions. Based on our metacommunity model, Eq. (1), we can now investigate the role of migration for metacommunities of weakly competing species and ask, how the balance of migration and stochastic extinctions shapes spatio-temporal abundance patterns in a species-rich metacommunity. In the following sections we will address this question for the two limiting scenarios of short-range migration and uniform migration between all patches (global migration). Our results from the fully symmetric case of indistinguishable species, Eq. (1), will provide insights that contribute significantly to the understanding of species-rich metacommunities with more general properties, which we will discuss in the final section.

### The migration rate needs to exceed a threshold to prevent global extinction

First, we consider migration on the smallest length-scale where individuals can migrate only between neighboring patches. This assumption has, for instance, been extensively applied in studies of expanding microbial biofilms (21–23, 47). To implement short-range migration in one dimension, we assume a one-dimensional lattice of patches, and set *λ*_*x,y*_ = (1*/*2)*λ* for all pairs of neighboring patches *x* and *y* and *λ*_*x,y*_ = 0 otherwise. The migration term in Eq. (1) then reduces to (1*/*2)*λ*(*N*_*x*+1,*i*_ + *N*_*x*−1,*i*_ − 2*N*_*x,i*_), where we furthermore assume periodic boundary conditions. First, we fix *r, K*, and *α* (with *α*≪1), and vary the migration rate *λ* for different numbers of species *S*.

When numerically solving the dynamics Eq. (1) with short-range migration (for details on the numerical solution, see *SI Appendix*, Sec. 1), we find that for zero and small migration rates *λ* all species eventually go extinct due to demographic fluctuations. In contrast, when *λ* exceeds a critical threshold value *λ*_*c*_, the average population size *N* = (*PS*)^−1^ _*x,i*_ *N*_*x,i*_ after the final time step of our numerical solution is finite and increases with *λ* (see circles in Fig. 2*A*). Here, species occasionally go extinct on individual patches, but are able to recolonize these patches eventually (Fig. 2*B*). Close above *λ*_*c*_, the species’ mean population sizes are infinitesimal small so that interactions between them should be negligible. Consistently, we find that the critical migration rate *λ*_*c*_ does not depend on the competition strength *α*, nor the number of species *S* (see Fig. 2*A*). Equations of the form of Eq. (1) without inter-species interactions (i.e. *α* = 0) are well-studied in the context of *directed percolation* [see (48, 49) for reviews] and have been broadly applied to emulate spreading processes such as forest fires and epidemics as well as range expansions in microbial biofilms (21–23, 47). In particular, for one-species metacommunities, the interplay of population growth, migration, and demographic fluctuations is known to show a non-equilibrium phase transition from a phase of zero population size (inactive phase) to a phase of finite population sizes (active phase) marking the directed percolation threshold. In our multispecies metacommunity, we thus recover the directed percolation threshold as the critical migration rate *λ*_*c*_, independent of the interaction strength *α* and the number of interacting species *S*.

**Fig. 2.**
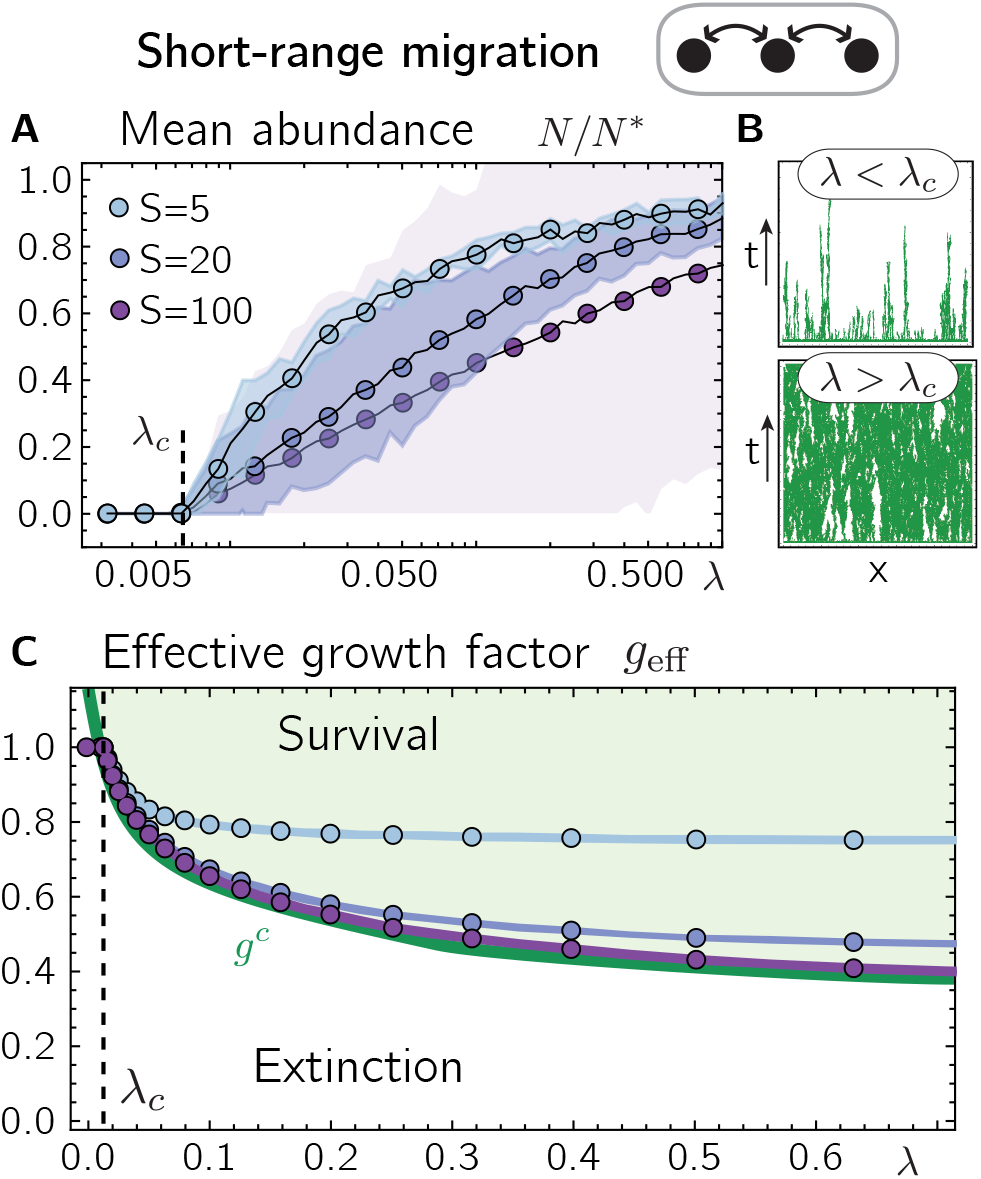
Migration-extinction balance and self-organization to the extinction threshold. **(A)** Numerical solutions of Eq. (1) for short-range migration show that for migration rates *λ* below *λ*_*c*_, all species go globally extinct whereas for larger migration rates the mean population size, *N* (circles), assumes non-zero values. Shaded areas denote standard deviations of patch-averaged abundances *N*_*i*_ across all species. **(B)** Spatio-temporal dynamics of a representative species for migration rates below and above the threshold *λ*_*c*_ for *S* = 5. Axes denote time *t* and the location (patch) *x*. Regions where the species is present and extinct are colored in green and white, respectively. **(C)** For migration rates *λ* larger than *λ*_*c*_, the mean effective growth factor 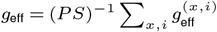 (circles) drops below one. The shaded areas denote the standard deviation of the patch-averaged effective growth factors *g*^(*i*)^ across all species (which is virtually zero). For increasing *S*, the effective growth factors asymptotically approach the single species threshold value *g*^*c*^ (green solid line). The green and white shaded areas indicate parameter regimes where the single species dynamics, Eq. (4), yields finite and zero population sizes, respectively. Parameter values are *r* = 0.3, *K* = 10, *α* = 0.1, *P* = 500. As initial condition we chose *N*_*x,i*_ = *K* for all patches and species with small random perturbations.

Comparing the patch-averaged population sizes of individual species, *N*_*i*_ = *P* ^−1^ Σ_*x*_ *N*_*x,i*_, after the last time step of our numerical solutions, we find that these can strongly differ across species, especially when the number of species is large (compare shaded areas in Fig. 2*A*). In particular, some species occupy only a very small fraction of patches. Some species also die out globally even for *λ > λ*_*c*_, which we attribute to the finite number of patches in our numerical solution.

### Species packing pushes growth rates towards extinction threshold

Motivated by the wide variation in population size across species, in the following we aim to better understand the metacommunity dynamics on the individ-ual species level. To investigate the dynamics of individual species, we first rewrite the deterministic growth dynamics of a species *i* on patch *x* [first term in Eq. (1)] as

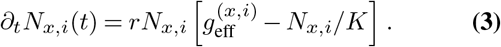

Here, we defined the *effective growth factor* 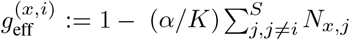 of species *i* on patch *x*, which can be understood as the ratio of the species’ growth rate in the presence of competing species and its growth rate in the absence of competing species (i.e. *r*). Depending on the degree to which inter-species competition suppresses population growth, the effective growth factor thus takes on values less than or equal to one. Eq. (3) suggests that—in the absence of migration and demographic fluctuations—a species’ population will grow and assume a finite population size precisely if its effective growth factor is larger than zero. Ignoring demographic fluctuations, this observation has been used to derive expressions for the abundance distributions in well-mixed species-rich communities with small constant immigration when inter-species interactions are randomly distributed (14, 16, 50).

In our numerical solutions of Eq. (1), we observe that the patch-averaged effective growth factors of a species, 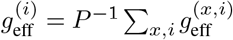, are virtually identical across different species (see Fig. 2*C*; see *SI Appendix*, Sec. 2 for their distribution). First, we find that for migration rates *λ* above the critical migration rate *λ*_*c*_, the patch-averaged effective growth factors 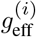 drop to values below one, consistent with the fact that there species coexist and thereby suppress each others growth through competition. Furthermore, when the number of species *S* increases, 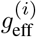 decreases until it eventually *saturates* at a positive finite value. Why does the patch-averaged effective growth factors saturate for large *S* and what is the role of this saturation for the spatio-temporal dynamics of the metacommunity?

To better understand the role of the effective growth fac-tor 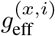 in the metacommunity’s dynamics, we substitute 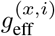 in Eq. (3) for a fixed parameter *g* and numerically solve the dynamics of the metacommunity for different choices of *g* and *λ* including demographic fluctuations and migration. Species are thus no longer coupled with each other and the dynamics of every species’ population size *N*_*x*_ on a patch *x* reduces to

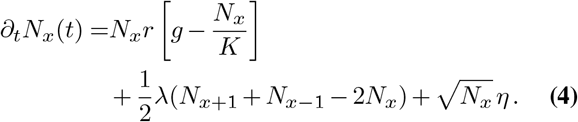

From directed percolation theory (48, 49), we expect Eq. (4) to feature a transition from a state of zero population size to a state of finite population sizes depending on the parameters *r, g, K*, and *λ*. Indeed, solving the one-species dynamics Eq. (4) numerically for different *g* and fixed *r, K*, and *λ*, we identify a threshold value for *g*, we denote *g*^*c*^ (see green line in Fig. 2*C*). When *g* is smaller than *g*^*c*^, the dynamics Eq. (4) eventually leads to stochastic extinction while for *g* larger than *g*^*c*^ the dynamics Eq. (4) leads to finite population sizes (see regimes ‘Survival’ and ‘Extinction’ in Fig. 2*C*, respectively). Interestingly, we find that this threshold value *g*^*c*^(*r, K, λ*), marks the asymptotic values of the patch-averaged effective growth factors 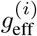 in the meta-community for large *S*:

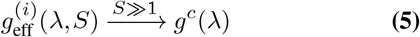

Hence, when the number of competing species is large, the patch-averaged effective growth factor of each species is pushed towards the critical threshold value, at which the species’ growth is just strong enough to balance stochastic extinctions. This suggests that every species operates close to its extinction threshold (percolation threshold). Importantly, this self-organization is not restricted to a particular choice of the migration rate or the remaining model parameters *r, K*, and *α*, but occurs for any *λ > λ*_*c*_ when the number of species *S* is large. In the *SI Appendix*, Sec. 3, we systematically increase the interaction strengths and argue that the observed self-organization towards the critical extinction threshold, Eq. (5), is present as long as the number of coexisting species at each patch (local diversity) is much greater than one, which is especially the case for weak species interactions.

From a physical perspective, we expect that close to a critical transition the characteristic time and length scales of the system’s dynamics diverge and the system’s observables obey scaling laws that are—to some extent—independent of model details, such as microscopic interaction assumptions (48, 49, 51). Indeed, when the number of species is large, our numerical solutions display spatio-temporal extinction patterns whose length and time scales extend to scales comparable to the system size and simulation time of our numerical solution, respectively (see Fig. 3*A,B*). More precisely, we find that the length 𝓁 of connected regions in which individual species are extinct is well-approximated by a power-law distribution (see Fig. 3*C*, purple circles). Similarly, the time *τ* between a species’ extinction on a patch and its successful recolonization of that patch from adjacent patches follows a power law distribution (see Fig. 3*D*, purple circles). Our above analyses indicate, that if the number of species in the metacommunity is large, each species’ dynamics follow the dynamics of a species in the absence of competitors for parameter values close to the percolation threshold. To further test this hypothesis, we numerically solved the single species dynamics, Eq. (4), for values of *g* close to *g*_*c*_ (for instance, when we choose 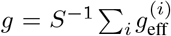 for *S* = 100). We find that the distribution of extinction lengths and times follow power-laws that are in excellent agreement with the distribution we found for the species-rich metacommunity (compare green and purple circles in Fig. 3*C* and *D*). Furthermore, these power-laws are well-described by exponents found in one-dimensional directed percolation of a single species (52, 53) (see dashed lines in Fig. 3*C* and *D*; for a more detailed discussion of the observed power-law exponents see *SI Appendix*, Sec. 4). Together, our results strongly suggest that in the species-rich metacommunity each species follows the dynamics of an isolated species close to its extinction threshold. While the effective growth factors averaged over patches are driven toward the threshold *g*^*c*^ for all species, the effective growth factors can differ between different patches (see *SI Appendix*, Sec. 2). Specifically, we find that species experience effective growth factors 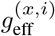 that lie below the threshold *g*^*c*^ on some patches. Our results thus show that species can survive effective growth factors below the threshold *g*^*c*^ on some patches as long as these patches are balanced by patches with effective growth rates above the extinction threshold, such that the patch-averaged effective growth factor of a species exceeds the extinction threshold.

**Fig. 3.**
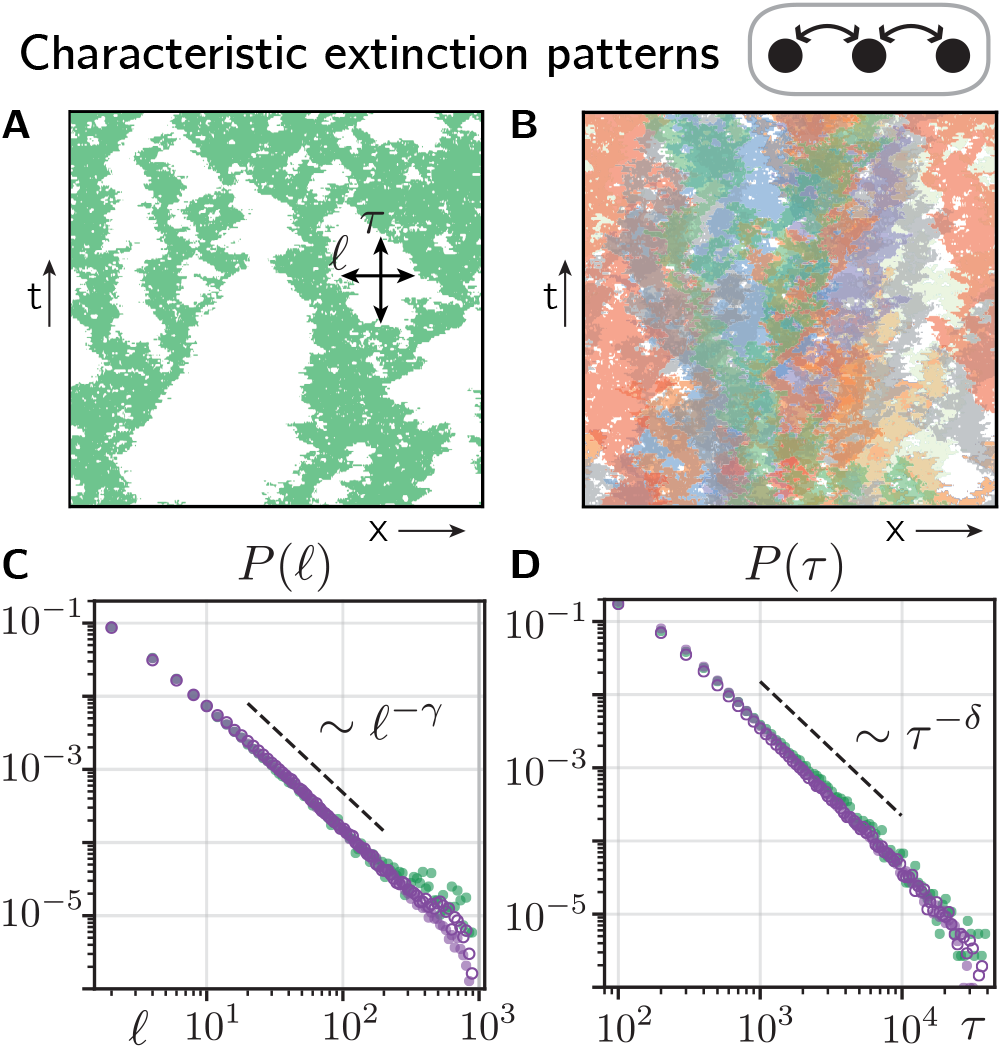
Characteristic spatio-temporal patterns in species-rich metacommunities. **(A)** For migration rates larger than *λ*_*c*_ (here, *λ* = 0.89), the spatio-temporal dynamics of a species in a species-rich metacommunity (*S* = 100) shows extinction patterns of various lengths 𝓁 and times *τ* that can range up to the system size and the time of our numerical solution, respectively. 200 time steps (generations) between generation 10^4^ and 5 × 10^4^ are shown. **(B)** Spatio-temporal dynamics of nine randomly picked species for parameters as in *A*. **(C)** and **(D)** show the distributions *P* (𝓁) and *P* (*τ*) of extinction lengths 𝓁 and times *τ* [indicated in *A*], respectively. Purple open and closed circles denote the distributions for *S* = 100 with small and large migration rates (*λ* = 0.32 and *λ* = 0.89, both larger than *λ*_*c*_), respectively, and green circles show the distribution for the single single-species dynamics, Eq. (4), at criticality *g* ≳ *g*^*c*^ (*g* is set to 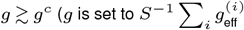 measured in a meta-community with *S* = 100). Dashed black lines indicate power-law distributions with exponents *γ* = −1.747 and δ= −1.840, respectively, as predicted from directed percolation theory. Parameter values are *r* = 0.3, *K* = 10, *α* = 0.1, *P* = 1000. As initial condition we choose *N*_*p,i*_ = *K* for all patches *x* ∈ {1,…*P*} and species *i* ∈ {1,…*S*} with small random perturbations.

As exemplified in various metacommunities, including microbial (20, 24) and plant communities (42, 54), different length scales of migration can confer very different statistical properties to an ecosystem, with important consequences on the evolutionary dynamics of the community. In the next section, we will therefore explore the question of whether and in what ways our results for short-range migration apply to larger length scales of migration.

### Species-rich metacommunities with global migration

So far, we have studied how metacommunity can maintain itself if we assume nearest neighbor migration. We now explore metacommunity dynamics under the opposite migration pattern of global, all-to-all migration. This migration pattern allows us to (i) check how sensitive our main results are to migration patterns and (ii) obtain concrete analytical results.

Specifically, we assume that all patches are connected through migration with a migration rate *λ*_*x,y*_ = *λ/P*. The migration term in Eq. (1) then reduces to 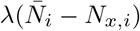 where 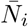 denotes the abundance of species *i* averaged over all *P* patches. For our analytical mean-field approach, we first express the interaction term Eq. (1) through the species-averaged abundance on a patch defined as 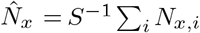 Then, by treating the mean-fields 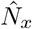 and 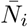 as deterministic mean-field parameters, we can map the dy-namics in Eq. (1) to the solvable problem of a Brownian par-ticle in a fixed potential. Since in our basic model, Eq. (1), all species are indistinguishable, the mean abundances 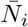 and 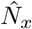 are equal in equilibrium (in the limit of an infinite number of species and patches). Finally, we can derive an analytic expression for the abundance distribution as a function of the mean species abundance 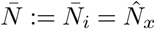 and the control pa-rameters *r, K, α, S*, and *λ* (for a detailed derivation see *SI Appendix*, Sec. 5). The abundance distribution is given by:

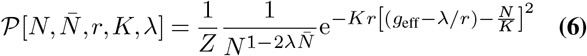

where *Z* denotes the normalization constant and we defined the mean-field effective growth factor 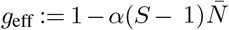. We can now solve for the mean abundance 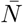 self-consistently by calculating the statistical mean abundance based on the distribution Eq. (6), ⟨*N* ⟩_P_, and demanding that 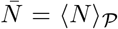. Eventually, this yields a closed form of the species abundance distribution. From the species abundance distribution, we can then calculate various equilibrium quantities such as the mean-field effective growth factor and the mean local diversity (see *SI Appendix*, Sec. 5 and 6). Depending on the choice of parameters, the abundance distribution 𝒫 approaches forms which have been commonly found in natural ecosystems (55–58) and mathematically derived from previous ecological models (57, 59–61). For instance, when the migration probability is small, i.e.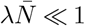, Eq. (6) follows the scaling 𝒫 [*N*] ∝*x*^*N*^ */N*, which is commonly referred to as Fisher log series and denotes one of the most widely used abundance distributions in ecology [see (62, 63) for reviews]. For larger migration rates, Eq. (6) suggests a Gaussian contribution with a maximum at *N* = *K*(*g*_eff_ −*λ/r*) and a variance *K/*(2*r*) (see *SI Appendix*, Sec. 5 for details).

Similar to short-range migration (compare Fig. 2*A*), we find that the mean abundance of all species undergoes a bifurcation at a critical migration rate *λ*_*c*_ from zero to non-zero values (see *SI Appendix*, Sec. 5). For *Kλ* ≪*Kr*, the critical migration rate can be approximated by

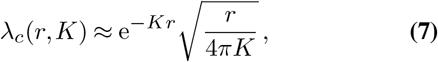

which is independent of the number of interacting species and the interaction parameter *α*. In the limiting case of *Kr Kλ* we obtain the approximation

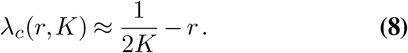

Both limiting behaviors, Eq. (7) and Eq. (8), are in very good agreement with numerical solutions of the metacommunity dynamics (see *SI Appendix*, Sec. 5). Moreover, when the number of species *S* increases, we find that the mean effective growth factor *g*_eff_ asymptotically approaches the single species threshold value *g*^*c*^(*λ*) (see solid lines in Fig. 4*A*; for the mean-field solution of *g*^*c*^ as well as its limiting behaviors for small and large migration rates, see *SI Appendix*, Sec. 6). Similar to short-range migration, this suggests that species in the species-rich metacommunity operate close to their extinction threshold.

**Fig. 4.**
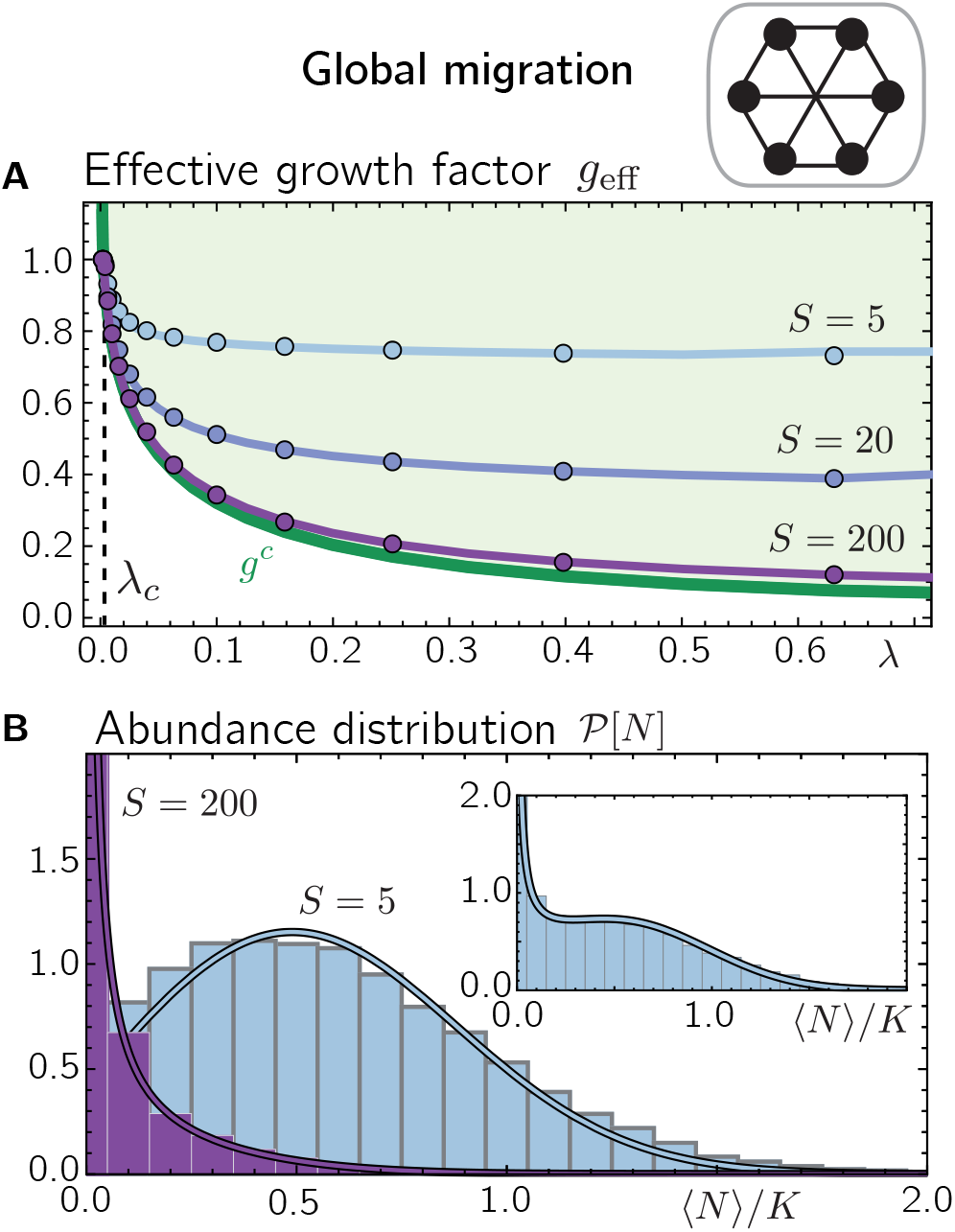
Mean-field approach is in good agreement with numerical simulations for global migration. **(A)** For migration rates *λ* larger than *λ*_*c*_, the mean effective growth factor *g*_eff_ drops below one, (circles and solid lines show numerical solutions and mean-field solutions for *g*_eff_, respectively). For large *S*, the mean effective growth factor asymptotically approaches the single species threshold value *g*^*c*^ (green solid line, calculated from mean-field theory). **(B)** Abundance distributions for *S* = 5 (light blue) and *S* = 200 (purple) for *λ* = 0.1 (main plot) and *λ* = 0.04 (inset). Histograms display the numerical solutions with global migration and solid lines show the corresponding mean-field solutions. Parameter values are *r* = 0.3, *K* = 10, *α* = 0.1. For the numerical solutions we choose the initial condition *N*_*p,i*_ = *K* for all patches *p* ∈ {1,…*P*} with *P* = 500 and species *i* ∈ {1,…*S*} with small random perturbations.

Treating 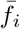 and 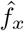 as deterministic mean-fields is based on the assumption that the number of patches *P* and the number of species per patch is large enough such that fluctuations in 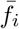 and 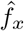 across species and patches, respectively, are negligible. While *P* can simply be chosen large in our numerical solutions, the diversity per patch depends on the model parameters including the competition strength *α*. When the diversity per patch is much larger than one, such as for weak competition (0 *< α* ≪ 1), as is the main focus of this work, we find very good agreement between our mean-field predictions and our numerical solutions (compare Fig. 4*A,B*). However, for larger *α*, especially *α* ≲ 1, the diversity per patch can drop to only one species and we observe deviations between our mean-field and numerical solutions (for a more detailed discussion of limitations of our mean-field theory see *SI Appendix*, Sec. 7).

### Variation in growth parameters drives the extinction of a part of the community

The proximity of species to extinction in a species-rich metacommunity allows several implications about the sensitivity of the metacommunity to perturbations. For instance, in a species-rich metacommunity, even a small variation in the migration rates or growth rates between species may lift the effective growth factors of some species below the extinction threshold and thereby lead to their global extinction. To investigate the effect of differences in the species’ growth dynamics and dispersal, we generalize Eq. (1) and consider the following dynamics in the metacommunity:

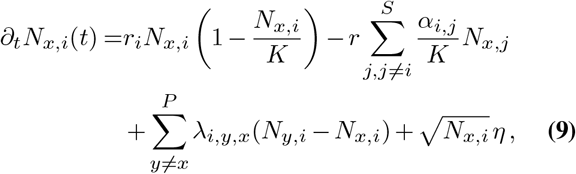

where fitness differences between species *i* are implemented by assuming differential growth rates *r*_*i*_. Furthermore, interactions between species *i* and *j*, represented by the coefficient *α*_*i,j*_, may differ, and species may have different migration rates *λ*_*i*_. For simplicity, the parameters *r*_*i*_, *λ*_*i*_, and *α*_*i,j*_ are drawn from normal distributions centered around *r, λ*, and *α*, with standard deviations *σ*_*r*_, *σ*_*λ*_, and *σ*_*α*_, respectively (negative migration rates are set to *λ*). Previous studies (11, 14, 16, 17, 50) have shown, that in a well-mixed ecosystem without demographic fluctuation and small differences in inter-species interactions 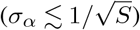, all species can coexist and the community approaches a unique stationary stable state. When inter-species interaction differ more strongly, a fraction of species will go extinct and the system shows multistability, which can lead to chaotic dynamics in metacommunities (17, 34).

With the generalized dynamics Eq. (9), the effective growth factor of a species *i* at the location *x* is given by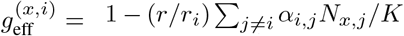. When we solve Eq. (9) numerically for short-range migration and relatively small parameter differences across species, in particular 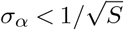, we find that the patch-averaged effective growth factors 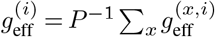 initially undergo a quick relaxation dy-namics followed by weak fluctuations around their steady states. Fig. 5*A* shows the patch-averaged effective growth factors 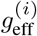 after the last time step of our numerical solution of Eq. (9) when only the competition strengths and migration rates vary between species (*σ*_*α*_, *σ*_*λ*_ *>* 0), and all species have equal fitness (i.e. *σ*_*r*_ = 0, *r*_*i*_ = *r*). First, we observe that compared to Fig. 2*C*, the patch-averaged effective growth factors 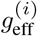 now vary more strongly between species. Specifically, species also assume patch-averaged effective growth factors that are relatively far above the critical threshold *g*^*c*^(*λ*). Other species assume patch-averaged effective growth factors below *g*^*c*^(*λ*) and die out eventually in the metacommunity (gray circles in Fig. 5*A*). Such global extinctions already occur for relatively small differences in interspecies interactions 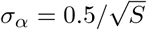, where coex-istence of all species is supposedly still a stable solution of the deterministic dynamics (11, 14, 16, 17, 50). Our results hence highlight the important role of demographic fluctuations in a species-rich metacommunity that is poised at the critical extinction threshold as it is in our case. Considering the surviving species, their patch-averaged effective growth factors (purple circles in Fig. 5*A*) cluster close to the critical extinction threshold *g*^*c*^(*λ*) with the majority of species being close to their (species-specific) extinction thresholds. When allowing differences in fitness and the interaction coefficients (i.e. *σ*_*r*_, *σ*_*α*_ *>* 0; *σ*_*λ*_ = 0), we observe a qualitatively similar behavior in the metacommunity (see Fig. 5*B*): While some species assume a patch-averaged effective growth factor relatively far beyond the threshold value *g*^*c*^(*r*) (plotted as a function of *r* in Fig. 5*B*), others assume a patch-averaged effective growth factor below *g*^*c*^(*r*), and eventually die out globally. The majority of surviving species assumes patch-averaged effective growth factors close to the threshold *g*^*c*^(*r*), suggesting that most species operate at their (species-specific) extinction thresholds. Solving Eq. (9) for global migration, we find a phenomenology similar to short-range migration (see *SI Appendix*, Sec. 8, for a more detailed discussion). In particular, species assume different patch-averaged effective growth factors 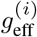 depending on the variances of differential growth rates, migration rates and interaction coefficients (i.e. *σ*_*r*_, *σ*_*λ*_, and *σ*_*α*_, respectively). For moderate *σ*_*α*_, species with 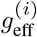 below the critical threshold *g*^*c*^ eventually go extinct globally, while the 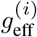 of the the surviving species cluster close above *g*^*c*^. Increasing *σ*_*α*_ leads to an increasing spread in patch-averaged growth factors 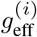 (*SI Appendix*, Sec. 8). For large *σ*_*α*_, we find that some species survive despite having a patch-averaged growth factor 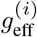 below threshold 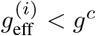 We hypothesize that this is related to the existence of multiple stable communities for large *σ*_*α*_, as sug-gested in (14, 17) (see *SI Appendix*, Sec. 8 for a more detailed discussion).

**Fig. 5.**
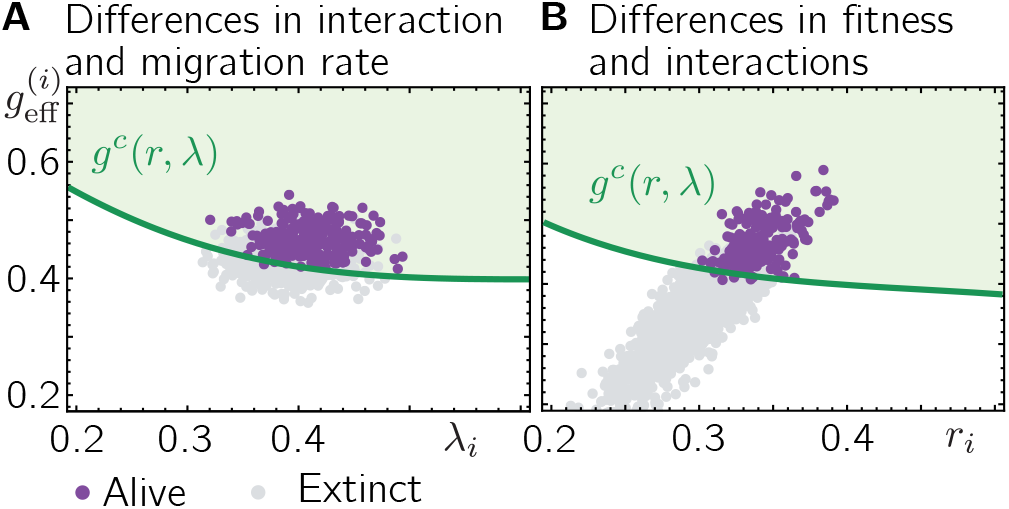
Variation in growth parameters leads to a loss of diversity in species-rich metacommunities. **(A)** Purple and gray circles denote the mean effective growth factors 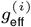 of species that are alive or have gone extinct at the end of our numerical solution, respectively. Interaction strengths and migration rates are drawn from normal distributions with mean *α* and *λ*, and standard deviations *σ*_*α*_ and *σ*_*λ*_, respectively (all growth rates *r*_*i*_ are set to *r*). The green line denotes the extinction threshold value for the growth factor *g*_*c*_(*λ*) [compare Fig. 2*A*]. When the number of competing species is large, the mean effective growth factors of the surviving species cluster close to the threshold value *g*_*c*_(*λ*). **(B)** Fitness *r*_*i*_ and interaction coefficients *α*_*i,j*_ are drawn from normal distributions with mean *r* and *α*, and standard deviations *σ*_*r*_ and *σ*_*α*_, respectively (all migration rates *λ*_*i*_ are set to *λ*). The green line denotes the extinction threshold value for the growth factor as a function of the growth rate *r*, i.e. *g*_*c*_(*r*). When the num-ber of competing species is large, the mean effective growth factors 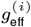 cluster close to the threshold value *g*_*c*_(*r*). The remaining parameter values are *r* = 0.3, *K* = 10, *λ* = 0.4, *α* = 0.1, *S* = 300, *P* = 2000; for *A*: *σ*_*λ*_ = 0.03,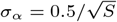; for *B*: *σ*_*r*_ = 0.03,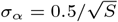. As initial condition we chose *N*_*x,i*_ = *K* for all patches and species with small random perturbations. *A* and *B* show the results for three independent draws of parameters and initial conditions.

## Discussion

In this study, we have explored the maintenance of biodiversity in closed metacommunities. In contrast to mainland-island models, long-term species survival requires that local species extinction is balanced by immigration from within the metacommunity. This leads to a minimal migration threshold, below which species invariably go extinct. Interestingly, even if migration rates exceed this threshold, species tend to self-organize close to an extinction threshold - the more so, the more species are added to the community. The qualitative agreement of our results on both limiting length scales for dispersal, i.e. short-range migration and global migration, suggests that this self-organization process is a general property of species-rich metacommunities and not restricted to certain length scales of migration.

In the case of short-range migration, we find that living at the edge of extinction generates fractal spatio-temporal dynamics, characteristic of a well-known non-equilibrium phase transition (directed percolation). Length and time scales of species extinction patterns can thus span the entire size and lifetime of the metacommunity, and their distributions follow power laws with exponents that are independent of system parameters such as growth and migration rates. In contrast to standard directed percolation, this behavior is not restricted to a single point in parameter space (e.g. a critical migration rate), but occurs whenever the number of species is large.

While empirical data indicates that spatially-averaged static observables, such as abundance distributions, follow rather general trends across different ecosystems (55–58) (57, 59– 61, 64, 65), our study gives also insights into dynamical properties of metacommunities. This allows to use much more specific spatio-temporal data to verify or falsify our model and to determine species’ proximity to global extinction.

We found that even small variations in the species’ growth, interaction, and migration rates lead to extinctions of a fraction of species where the ability to survive can be characterized by a species’ patch-averaged effective growth factor. This has several implications for the manipulation and preservation of species-rich metacommunities in the view of a changing environment. For instance, environmental perturbations that result in slightly different growth parameters among species (even if transient) can cause a large number of species to go extinct that previously coexisted near their extinction thresholds. In addition, species that are prone to extinction can be saved by selectively increasing their fitness or migration rate on some patches so that their patch-averaged effective growth factor falls above the critical extinction threshold. In the course of evolution, we hypothesize that the need for a species to overcome a non-zero critical growth factor to survive may have important consequences for its fixation probability and thus the evolution of species-rich metacommunities.

By considering weak inter-species competition among species where demographic fluctuations dominate the dy-namics, our work provides a natural counterpart to several previous studies. These include work on metacommunities with strong fluctuations among patches due to the species’ interactions (33, 34) as well as neutral models (8, 65, 66), which–even in structured metacommunities (57, 67–69)–rely on a continual immigration from a species pool. Moreover, our study suggests generalizations of previous work on mainland-island models (14, 16, 17, 50) towards closed metacommunities with demographic fluctuations, which we expect to generally feature migration thresholds.

## ACKNOWLEDGEMENTS

We thank current and former members of the Hallatschek lab, especially Stephen Martis and Takashi Okada, for helpful comments and discussions, and Daniel S. Fisher (Stanford University) for stimulating discussions on complex metacommunities. Research reported in this publication was supported by a National Science Foundation CAREER Award (1555330) and by the German Research Foundation through grant 445916943 (to J. D.).

## Supplementary Note 1: Numerical solution of the metacommunity dynamics

As detailed in the main text, the metacommunity is assumed to follow the dynamics:

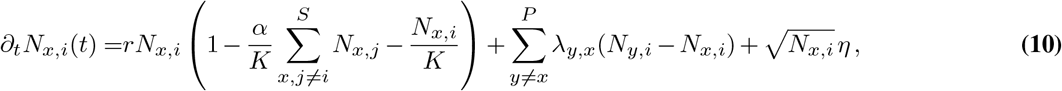

where *N*_*x,i*_ denotes the abundance of species *i* ∈{1, …*S*} on the patch *x* ∈{1, …, *P*}. The first term denotes population growth and interactions with other species, the second term denotes migration between patches, and the third term accounts for demographic fluctuations where *η* is uncorrelated noise with zero mean and variance *ω*^2^. By rescaling the growth rate and the migration rate with the rate *ω*, we measure time in units *ω*^−1^ and can set *ω* = 1 in the following. Unless noted otherwise, for Fig. 2-5 we fixed the growth rate (*r* = 0.3), the competition strength (*α* = 0.1), the carrying capacity (*K* = 10), and solved the dynamics for short-range and global migration for various migration rates *λ* and different numbers *S* of initially coexisting species based on the following Euler forward scheme [all calculations were performed in `Python` (70) and the results were evaluated using `Mathematica` (71).]. For each time step Δ*t*, we first calculate the update of the population size for each species on each patch given by the growth and migration dynamics [first two terms in Eq. (10), respectively]. Short-range migration is implemented on a one-dimensional lattice with *P* sites where we set *λ*_*x,y*_ = (1*/*2)*λ* for all pairs of neighboring patches *x* and *y* and *λ*_*x,y*_ = 0 otherwise (we further assume periodic boundary conditions, i.e. *N*_*P* +*x,i*_ = *N*_*x,i*_). For global migration, we set *λ*_*x,y*_ = *λ/P* so that a patch is colonized at a rate 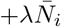, with 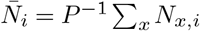. The contributions from growth and migration are calculated and updated separately in order to avoid unphysical scenarios, e.g. that an unoccupied patch acts as a source of migration. After updating the deterministic abundance of each species on each patch, demographic fluctuations [last term in Eq. (10)] are added by sampling from a Poisson distribution with the mean being the deterministic abundance. Interpreting Eq. (10) in the Itô sense (72), the Euler forward update for the demographic fluctuations are then incorporated by adding 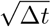(`Poisson` [*N*_*x,i*_] − *N*_*x,i*_) to the deterministic abundances, where `Poisson`[*N*_*x,i*_] is a sample from a Poisson distribution with mean *N*_*x,i*_. The implementation of demographic fluctuations through a Poisson process guarantees that their variance is given by *N*_*x,i*_ (72), consistent with Eq. (10). As initial condition we choose *N*_*p,i*_ = *K* for all patches *p* ∈{1, …*P*} and species *i* ∈{1, …*S*} with small random fluctuations. Even beyond the onset of finite mean population sizes and in the totally symmetric case, Eq. (10), where all species have identical growth, interaction, and migration rates, we observe that a fraction of species goes extinct globally in our numerical solution. We argue that these global extinctions are due to the finite size of our system (finite number of patches *P*) and should vanish in the thermodynamic limit *P* → ∞ For the numerical solution of the more general metacommunity given by Eq.9 in the main text we employ an analogous Euler forward scheme as above where before every numerical solution, the interaction strenghts *α*_*i,j*_, the migration rates *λ*_*i*_ and the differential growth rates *r*_*i*_ are drawn from normal distributions centered around *α, λ*, and *r*, with standard deviations *σ*_*α*_, *σ*_*λ*_, and *σ*_*r*_, respectively (negative migration rates are set to *λ*). For our numerical solutions, the time steps Δ*t* are adapted to values between 0.2 and 1 (when the time step is chosen too large, the deterministic dynamics can generate negative population sizes, especially when the number of species *S* is large). The last time step in our numerical solutions ranges between 20000 (Fig.2,4,5) and 50000 (Fig.3), measured in units *ω*^−1^. The Python code developed for this study is available at https://github.com/Hallatscheklab/Self-Consistent-Metapopulations.

## Supplementary Note 2: Distribution of effective growth factors

In the totally symmetric metacommunity, i.e. when the growth rate *r*, carrying capacity *K*, inter-species competition strength *α*, and migration rate *λ* are chosen identical for all species, the effective growth factor for species *i* on the patch *x* is defined as

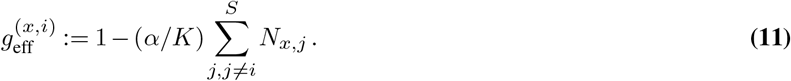

For the parameters chosen in the main text, particularly 0 *< α* ≪1 (e.g. see Fig.2C), the patch-averaged growth factors, 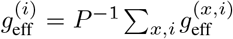, are virtually identical for all species. Fig. S1*A* displays the effective growth factors *g*^(*x,i*)^ for all patches for a representative species for different migration rates and numbers of species (*S* = 5 and *S* = 100). Furthermore, Fig. S1*B* and *C* show the distributions of effective growth factors *g*^(*x,i*)^ for different species and patches at certain migration rates close to the extinction threshold (*B*) and farther beyond the extinction threshold (*C*). In general, we find that the distribution of effective growth factors (including the mean) are very similar across different species. Furthermore, the effective growth factors of a species can assume values below the threshold value *g*^*c*^ on individual patches without causing the species’ global extinction (Fig. S1*A*). Our results thus suggest that effective growth factors that lie below the extinction threshold on some patches can be balanced by patches with larger effective growth rates and that species can persist in the metacommunity as long as their patch-averaged effective growth factor 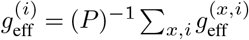 exceeds the extinction threshold *g*^*c*^.

## Supplementary Note 3: Increasing the competition strength can lead to spatial demixing

As discussed in the main text, in this study we focus on scenarios of weak inter-species competitions with 0 *< α* ≪ 1, so that multiple species can coexist on the same patch. For 0 *< α <* 1, the deterministic dynamics of Eq. (10) possesses a stable steady state solution where all species coexist on all patches. In contrast, when *α >* 1, the deterministic dynamics on an isolated patch approaches a steady state with only one species present (43, 45) (referred to as competitive exclusion). This suggests that in a metacommunity with *α >* 1, species may exclude each other on the same patch and occupy different sets of patches (provided migration is weak enough to allow heterogeneity between patches), as assumed in previous studies (58, 60, 73). In this section we discuss our observation that in the presence of demographic fluctuations a metacommunity can exhibit exclusion of species on the same patch even for *α <* 1. To this end, we increase the competition strength *α* from zero (i.e. no interaction) to values up to *α >* 1. First, we observe that the mean local diversity generally increases from zero at the threshold *λ* = *λ*_*c*_ to larger values with increasing migration rate *λ* (red symbols in Fig. S2*A*). Furthermore, the mean local diversity decreases with increasing *α*; for large *α* it falls to values close to and even below one even far beyond the threshold, i.e. for *λ* ≨ *λ*_*c*_ (e.g. see red triangles for *α* = 0.9 in Fig. S2*A*). The respective kymographs show that in these parameter regimes, species spatially demix and do not interact on most patches (Fig. S2*B*). Moreover, in these regimes of spatial demixing the mean abundance of all species follows the abundance of a single species in the absence of interactions. This is seen when plotting the mean abundance of species as a function of the migration rate: the bifurcation of the mean abundances indicates a transition between a regime where species demix and behave as if isolated and a regime where species coexist on patches (see squares in Fig. S2*C*).

In the regime where the local diversity is approximately one and species demix, they hardly interact on patches and the definition of an effective growth factor to characterize suppression of population growth by other species is no longer meaningful. We see that in these regimes, the measured patch-averaged growth factors 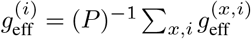 significantly differs between different species and that 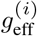 of some species, as well as the the mean effective growth factor 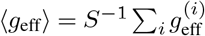, can fall below the threshold value *g*^*c*^ (Fig. S2*D*)

We think that this demixing of species for *α <* 1, where the deterministic dynamics alone would yield a spatially mixed metacommunity, poses an interesting question for future studies. As discussed in the main text, in the present work we are, however, particularly interested in scenarios where typically many more than one species coexist on the same patch (i.e. no species demixing) and inter-species competitions are weak 0 *< α* ≪ 1.

## Supplementary Note 4: Characteristic power-law exponents in species extinction patterns

Directed percolation denotes a class of non-equilibrium processes, that describe the spreading of non-conserved agent through a lattice (48). Implementations of directed percolation, such as the contact process as well as the Domany-Kinzel cellular automaton, have frequently been applied to model various spreading processes including forest fires, epidemics (for reviews see (48, 49)) and more recently range expansions of microbial biofilms (23, 47). A continuum description of directed percolation in *d* dimensions is given by the stochastic differential equation

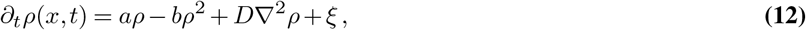

where *ρ*(*x, t*) ≥ 0 is the coarse grained density of the spreading agent (particle density) with space and time variable *x* and *t*, respectively, *a* and *b* are phenomenological parameters, ∇^2^ denotes the diffusion operators in *d* dimensions with diffusion constant *D*, and *ξ*(*x, t*) denotes a density-dependent Gaussian noise field with the correlations:

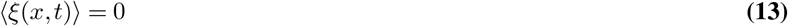

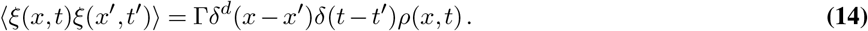

With these definitions, the functional form of Eq. (12) is the same as in Eq.4 in the main text with the discretized diffusion operator ∇^2^ → 0.5[*ρ*(*x* − Δ*x*) + *ρ*(*x* + Δ*x*) − 2*ρ*(*x, t*)] for a one-dimensional lattice with lattice constant Δ*x*. Importantly, the dynamics Eq. (12) shows a non-equilibrium phase transition (as function of the parameters *a*, Γ, and *D*) from an inactive phase, characterized by a density that decays exponentially with time, to an active state, where the density reaches a finite non-zero value. This transition is commonly termed *directed percolation threshold*. Combining field theoretic approaches with numerical solutions of Eq. (12), it was shown that, similarly to a equilibrium phase transition, the critical threshold between the inactive and the active phase is characterized by universal scaling laws for the macroscopic observables including the density. In particular, for a finite lattice of extension *L* the density *ρ* at the percolation threshold scales as

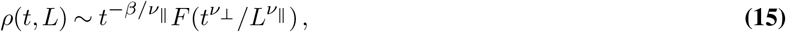

where *F* is some function that only depends on the variable combination 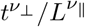. Here, *ν*_⊥_, *ν*_*∥*_, and *β* denote the characteristic scaling exponents for directed percolation. Following (52, 53) we can use the scaling Eq. (15) to derive the scaling of the distributions 𝒫 for the extensions 𝓁 and *τ* of connected unoccupied patches (voids) in space and time, respectively. Assuming the power-law behavior 𝒫 [𝓁] 𝓁^−*γ*^ we can write the mean size of connected voids in space within a space interval *L* in the stationary state as

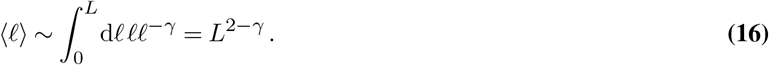

Moreover, from Eq. (15) and the assumption that *ρ* is finite for large *t*, we can derive 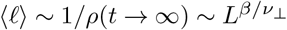. Hence, *γ* = 2 −*β/ν*_⊥_. Similarly, assuming the power-law behavior 𝒫 [*τ*] ∼*τ* ^−*δ*^ we can write the mean size of connected voids in time within a time interval *t* for a large enough system as

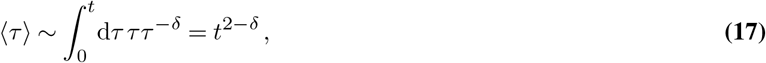

which, together with the scaling ⟨*τ* ⟩ ∼ 1*/ρ*(*t, L* → ∞) ∼ *t*^*β/ν*^_∥_ imposed by Eq. (15), yieldsδ= 2 − *β/ν*. With the values for *β, ν*_∥_, and *ν*_⊥_ found for one-dimensional directed percolation (48), we obtain *γ* ≈ 1.747 andδ≈ 1.840, which is in good agreement with the power laws found for 𝒫[𝓁] and 𝒫[*τ*] in the species-rich metacommunity (see Fig.3*C,D*).

## Supplementary Note 5: Self-consistent mean-field approach for global migration

In the following we will discuss our analytical mean-field approach to species-rich metacommunities with global migration that allows us to calculate static quantities such as the mean population size of species, the abundance distribution and the mean effective growth factor. For convenience, we describe our analysis based on the dynamics for the relative abundances *f*_*x,i*_ = *N*_*x,i*_*/K*, given by

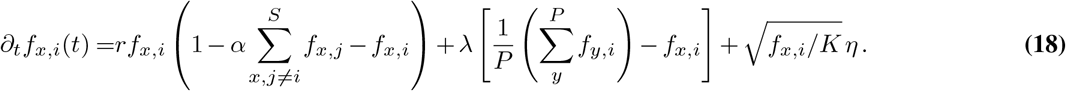

We introduce the mean fields 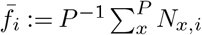 and 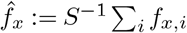, where 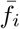 and 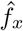 denote the averages of the relative abundance *f*_*x,i*_ taken over patches and species, respectively. In the following we assume that the number of patches *P* and the number of coexisting species on each patch are large, and that all species are statistically identical [with identical growth rates, carrying capacities, interactions and migration rates, as in Eq. (18)]. Under this assumption, we estimate the sums over different patches and species in Eq. (18) through the mean field expressions 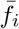 and 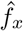 and treat these mean fields as deterministic parameters (for a more detailed discussion of this approximation see section 7). The dynamics of every species on every patches can then be expressed as

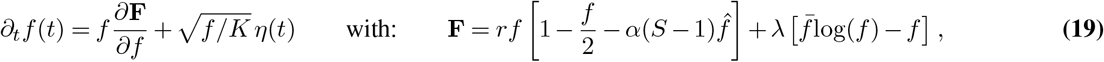

where we omitted the species and patch index since all species on all patches are statistically identical. The representation Eq. (19) admits an analytical equilibrium distribution in terms of a Gibbs measure (74–76). To see this, it is convenient to introduce the variable 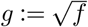. Using Itô’s lemma, we can rewrite Eq. (19) in terms of *g* as

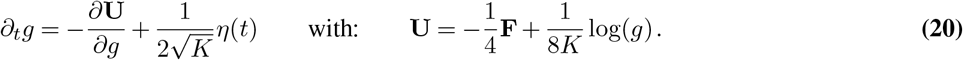

This dynamics for *g* can be reinterpreted as the overdamped dynamics of a particle in a potential **U** with diffusion constant 1*/*(4*K*). The equilibrium distribution 𝒫[*g*] for *g* is then given through the Gibbs measure

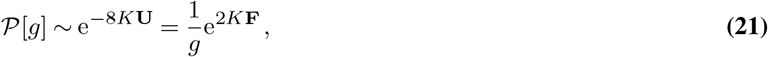

which is equivalent to a Boltzmann distribution with an “energy” given by **F**.

In terms of the relative abundance *f*, we have 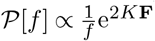, which can be expressed more conveniently as

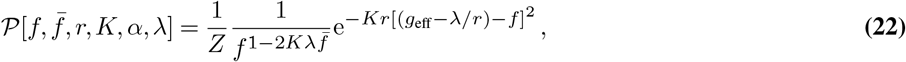

where we defined the mean-field effective growth factor 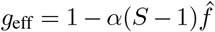 and *Z* denotes the normalization constant. In terms of the abundance *N* = *Kf*, the distribution can be written as

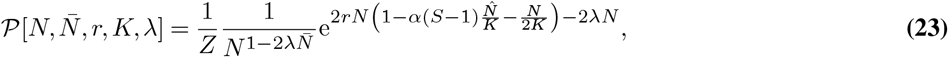

with respective normalization *Z*. Eq. (23). Eq. (22) is the result Eq. 6 in the main text with 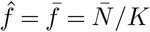. While we treated the mean fields 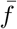 and 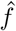 as deterministic parameters, in order for our analysis to be self-consistent they have to be equal and also equal the actual statistical mean of *f*, which can be calculated from the distribution Eq. (22). Introducing a Lagrange multiplier+*ϵf /*2*K* into the function F, we can take the derivative of log(*Z*) w.r.t to *ϵ*, take the limit *ϵ* → 0, and thereby obtain the mean abundance 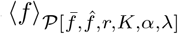. Self-consistency then requires:

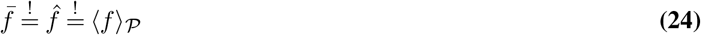

Fig. S3*A* shows the calculated mean ⟨*f*⟩ _𝒫_ as a function of 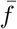 (where for specificity *r* = 0.3, *K* = 10, *α* = 0.1, and 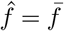 due to self-consistency). All calculations were performed using `Mathematica` (71). Varying the migration rate *λ* we find that for small *λ* the only solution to the self-consistency condition, Eq. (24), is given by 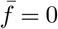. Increasing *λ* above a critical value *λ*_*c*_, the solution 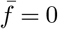 is no longer stable; however, there appears a second solution with non-zero 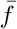, which is linearly stable and increases with *λ* [see Fig. S3*A*]. Thus, *λ*_*c*_ marks a bifurcation from zero to non-zero mean abundances. Expanding the calculated mean 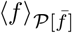 to first order in 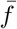, yields the condition for the onset of finite mean abundance:

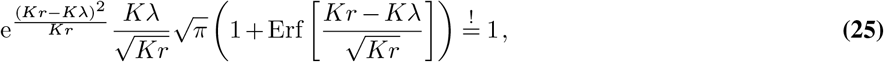

where Erf[·] denotes the Error-function (incomplete Gaussian integral). Note that the onset of finite mean abundances, Eq. (25), does not depend on the interaction strength *α* nor the number of interacting species *S*. This is consistent with the expectation that at the onset of finite mean abundances, interactions between species on a patch should be negligible. As a consequence, Eq. (25) equally describes the onset of non-zero mean abundances in a one-species metacommunity. The onset of non-zero mean abundances in a one-species metacommunity has been extensively studied in the context of percolation theory (48, 49), where it is referred to as percolation threshold.

### Critical migration rate

For the limiting case *Kλ* ≪*Kr*, we expand the condition Eq. (25) up to first order in *Kλ/*(*Kr*) and solved for *λ*, which yields the critical migration rate *λ*_*c*_:

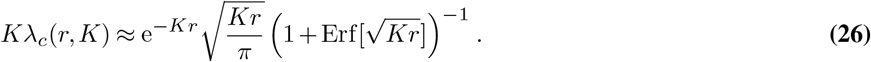

When furthermore *Kλ* ≪ 1, the growth rate must be correspondingly large so that we can set 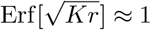. This yields 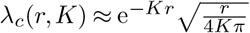, which is Eq. 7 in the main text. On the other hand, in the limiting case *Kλ* ≫ *Kr* we can expand the condition Eq. (25) to leading order of large *Kλ/*(*Kr*), and solve for *λ*. This yields the approximation (Eq. 8 in the main text):

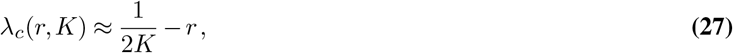

Hence, we find that for infinitesimally small growth rates (i.e. *r* →0), the critical migration rate approaches *λ*_*c*_(*r, K*) = 1*/*(2*K*). To explore the extreme case *r* = 0, we set the growth rate *r* equal to zero in the dynamics Eq. (19) and the resulting distribution Eq. (22). Calculating the statistical mean ⟨*f* _𝒫_⟩, we find that any mean abundance is marginally stable and fulfills the self-consistency equation Eq. (24). This suggests that fluctuations will ultimately (albeit on very long time scales) drive the metacommunity into the only absorbing state of the metacommunity *N*_*x,i*_ = 0. We comment that in the extreme case *r* = 0, the Langevin equation Eq. (10) resembles the well-studied voter model (48, 77, 78). Analogous to the voter model, which has two absorbing states (fully occupied and empty) and can show coexistence of these two states in dimensions larger than two, we expect that for *r* = 0 our metacommunity will eventually run in its only absorbing state *N*_*x,i*_ = 0. Both limiting behaviors of the critical migration rate *λ*_*c*_, at *Kλ/*(*Kr*) ≪ 1 and *Kr/*(*Kλ*) ≪ 1, are in very good agreement with respective numerical solutions of the Langevin equation Eq. (18) (see Fig. S4).

### Abundance distributions

Beyond the onset of finite mean abundances, we can solve for the mean abundance 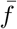 that satisfies the self-consistency condition Eq. (24) numerically [see Fig. S3*B*]. Eventually, substituting this numerical solution for 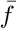 into Eq. (22) yields the equilibrium abundance distribution 𝒫 as a function of *r, K, α*, and the migration rate *λ*. For the parameters used in the main text, including Fig.4, the bifurcation from zero to finite mean abundances [see Fig. S3*B*] as well as the shape of the abundance distributions calculated by our mean-field approach (see Fig.4) are in very well agreement with explicit numerical solutions of the full, species-rich dynamics Eq. (18). In section 7 we discuss limitations of the presented mean-field theory and deviations between the mean-field prediction and numerical solutions of the metacommunity dynamics that appear when the mean number of species per patch is small or the abundance distribution is very broad.

### Characteristic contributions of the mean-field abundance distribution

Above the onset of finite population sizes (*λ> λ*_*c*_), the abundance distribution derived from our self-consistent mean field approach, Eq. (22), is governed by different contributions, depending on the choice of parameters: When 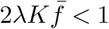, the exponent of *f* in Eq. (22) is negative and the abundance distribution Eq. (22) diverges at zero abundance. The mean relative frequency 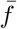 is limited by *f* ^*^ := *N* ^*^*/K*, where *N* ^*^ = *K/*[1 + *α*(*S* −1)] denotes the stationary uniform solution of Eq. (10) (see Eq.2 in main text). Since *f* ^*^ ∼ (*αS*)^−1^, the abundance distribution Eq. (22) shows a divergence at zero abundance whenever *S* is large enough, in particular when *S* ≳2*Kλ/α*. When the probability of migration is small 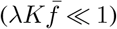 and for *f* ≪1 (i.e. abundances *N* ≪*K*), the distribution of the relative abundance *f* follows the scaling

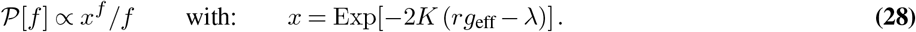

The form of the abundance distributions 𝒫 [*f*] ∝ *x*^*f*^ */f* is well-known in ecology literature as Fisher log series (79), which denotes one of the most widely used abundance distributions in ecology and has been recovered in a variety of ecological systems [see (62, 63) for reviews]. Furthermore, the Fisher log series has been derived on mathematical grounds for neutral ecosystems (i.e. where two individuals of the same or different species compete equally) with static immigration as limiting cases for small immigration rates [(59, 80), also see (81) for a review of analytic derivations of abundance distributions for neutral ecosystems]. Thus, our study shows that ecosystems that may exhibit static abundance distributions of neutral ecosystems may in fact underlie quite different species interactions.

For larger relative abundances (*f*∼1), the exponential term in Eq. (22) suggests a local maximum of the abundance distribution characterized by a Gaussian distribution with mean *g* _eff_ −*λ/r* and a variance 1*/*(2*Kr*). When the number of immigrants is small, 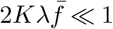, and growth is more likely than migration of an individual, *λ* ≪*rg*_eff_ the abundance distribution is bimodal, with a contribution for stochastic extinctions at *f* = 0 and a contribution at *f* ≈*g* _eff_ denoting occupied patches. In contrast, farther beyond the onset of finite abundances, 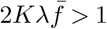, the abundance distribution is unimodal, and can be approximated by a Gaussian as mentioned above.

### Mean local diversity

Another interesting quantity in ecology is the diversity on a single patch (local diversity). The probability of a species on a patch to be extinct, **P**_0_, is the integral over the mean abundance distribution 𝒫 [*N*], Eq. (23), from zero to one. The mean number of species present on a patch is then given by *d*_1_ = *S*(1−**P**_0_). We can find a feasible analytical expression when we assume that the carrying capacity is large (*K* ≫1) and 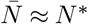. Then, the abundance distribution Eq. (23) can be approximated by

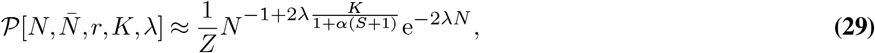

where *Z* denotes the respective normalization constant; we further assumed *S* ≫1 so that *N* ^*^*α*(*S*−1) ≈ *K*. The integral of Eq. (29) from zero to one can be easily calculated and division by the integral of Eq. (29) from zero to infinity eventually yields an approximation for **P**_0_. The respective approximation for *d*_l_ is then given by

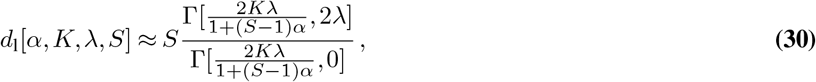

where Γ[·, ·] denotes the *Euler Gamma function* defined as 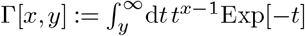. In the limit of large *S* this suggests that the mean local diversity approaches a finite value:

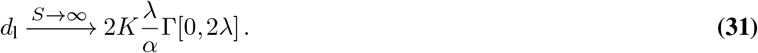

For the parameters chosen in Fig. S5A, we find reasonably good agreement between the estimate Eq. (30) (dashed lines) and the measured mean local diversity in our numerical solution (symbols and solid lines).

## Supplementary Note 6: Critical growth factor and single species percolation threshold

Following the mean-field analysis detailed in section 5, calculating the critical growth factor *g*^*c*^(*r, K, λ*) for global migration is straightforward. The effective one-species dynamics, Eq. 4 in the main text, in terms of the relative abundance *f*_*x*_ = *N*_*x*_*/K* for global migration can be written as

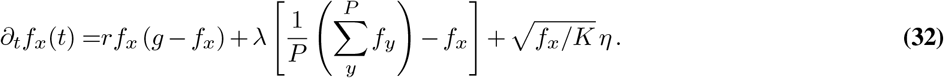

In analogy to section 5, we express the mean over patches through the mean field variable 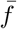, which is then treated as a deterministic parameter. We can then write Eq. (32) as

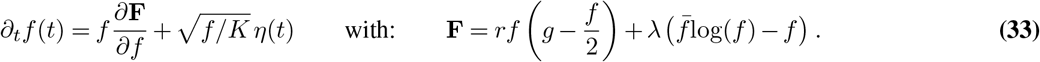

This equation has the same form as Eq. 20, allowing us to write the equilibrium distribution of ⟨*f*⟩ as a Gibbs measure. From the equilibrium distribution we then obtain an analytic expression for the mean relative abundance ⟨*f*⟩ as a function of the mean field.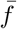. Imposing self-consistency, i.e.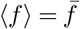, eventually yields an expression for the onset of non-zero population size given by

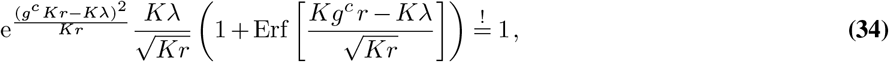

where *g*^*c*^ denotes the critical growth factor. Eq. (34) can be solved for *g*^*c*^ numerically [see Fig.4*A*], so that the critical growth factor *g*^*c*^ will be a function of the growth rate *r*, the carrying capacity *K*, and the migration rate *λ*.

In the limiting case *Kλ* ≪ *Kr* we can expand Eq. (34) up to first order in *Kλ/*(*Kr*) and find the approximation for *g*^*c*^:

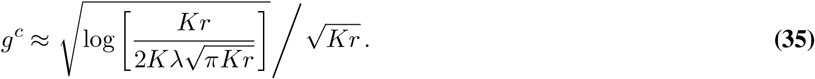

To obtain an approximation of *g*^*c*^ in the limit of large migration rates, we assume that *g*^*c*^*Kr* ≪*Kλ*, and expand Eq. (34) to first order in *g*^*c*^*Kr/*(*Kλ*). In this limit, the critical growth factor *g*^*c*^ can be approximated by:

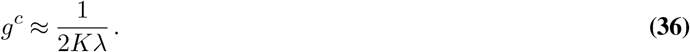

## Supplementary Note 7: Limitations of the analytic mean-field approach

As described in section 5, our mean-field approach is based on the assumption that we can express the sums over species and patches in Eq. (18) by their deterministic mean-field values 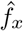 and 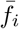, respectively. Substituting the sum over patches for the mean-field value 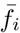 can be justified by choosing the number of patches *P* in the numerical solutions sufficiently large. On the other hand, if we express the sum of species on a patch by the deterministic mean-field 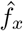, we implicitly assume that the typical number of coexisting species on a patch (local diversity) is sufficiently large. Specifically, the local diversity has to be large enough so that all species on a patch sample a major contribution of the abundance distribution. In the following, we give examples where the mean-field prediction starts to deviate from the numerical solutions as a result of low local diversities.

### Mean-field theory deviates from numerical solutions when competition between species increases

The deterministic steady state solution *N* ^*^ = *K/*[1 + *α*(*S*−1)] suggests that the number of species that can coexist (i.e. have an abundance *N*_*i*_ ≥1) on a patch is bounded by *S* ≤ (*K*−1 + *α*)*/α*. Moreover, for *α* ≥ 1, the coexistence state becomes deterministically unstable and an isolated patch will eventually approach a state where only a single species is present (45) (competitive exclusion). While in our study we focus on the scenario of competitions that are so weak that the local diversity is on average much larger than one, in the following we compare scenarios of stronger competitions–and consequently low local diversity–with our mean-field analysis. To this end, we increase *α* from zero (i.e. no interaction) to *α* ≳ 1 and compare our numerical solutions of the explicit multi-species dynamics, Eq. (18) (see section 1), with the predictions from our mean-field theory, section 5.

#### Bifurcation of the mean abundance

Fig. S5*A* shows the mean local and the global diversity (red and blue symbols, respectively) as a function of the migration rate for different choices of inter-species competition strength *α* and initial species *S* in the metacommunity. Close to the onset of non-zero mean abundance (i.e. *λ*≳ *λ*_*c*_) the mean local diversity increases from zero to larger values with increasing migration rate. Furthermore, the mean local diversity decreases with increasing *α* and may, for large *α*, drop to values around one even for migration rates far beyond the onset of non-zero abundance. Especially when the mean local diversity remains small (≈1) for migration rate beyond the onset *λ*_*c*_, we find deviations between the mean abundance in our numerical solutions (symbols in Fig. S5*B*) and our mean-field predictions (blue and purple lines Fig. S5*B*). Similar to the case of short-range migration (see section 3), we argue that in these parameter regimes species hardly interact with each other on patches but behave like isolated species that occupy only certain subsets of all patches of the metacommunity. In accordance with this conjecture, the mean abundance of our numerical solutions in these regimes approximately follows the mean-field prediction for the mean abundance in the absence of interactions, i.e. *α* = 0 (black dashed lines in Fig. S5*B*). Generally, it appears that the metacommunity assumes states where species are demixed (i.e. the mean local diversity is approximately one) or where multiple species occupy the same patches (i.e. mean local diversity much larger than one), depending on which state yields the larger total abundance for the metacommunity. This observation is reminiscent of our results with short-range migration (section 3). We think that the observed demixing of species with short-range and global migration, even for *α <* 1, poses an interesting question for future research. In this present study, however, we focus on weak inter-species competition 0 *< α* ≪1, in which case the local diversity is typically larger than one.

#### Effective growth factors

Similarly to short-range migration (section 3), we argue that in regimes where species demix and hardly interact on patches, the definition of an effective growth factor may no longer be meaningful to measure the suppression of population growth due to inter-species competition. As for short-range migration, we find that when the mean local diversity drops to values close to one, the patch-averaged effective growth factor differs more between species and the mean effective growth factor can fall below the critical threshold value *g*^*c*^ (symbols in Fig. S5*C* show the species and patch-averaged effective growth factors).

#### Abundance distributions

In accordance with the observations above, we find that the abundance distributions in our numerical solutions (symbols in Fig. S5*D*) show very well agreement with the mean-field predictions [blue and purple lines in Fig. S5*D*] when the mean local diversity is much larger than one. In regimes of small mean local diversities (≈1), the numerical solutions are better described by a mean-field prediction where species do not interact (see black dashed lines in Fig. S5*D*).

In summary, our mean-field approach yields a very good estimate for the metacommunity equilibrium when the local diversity is much larger than one. In particular, we find very good agreement for weak competition strengths (0 *< α* ≪1), which is the main focus of this work. We find that in parameter regimes with stronger inter-species competitions *α* (however, already for *α <* 1) and intermediate migration rates *λ*, species can demix due to demographic fluctuations, which leads to deviations from our mean-field predictions. We observe spatial demixing due to demographic fluctuations for both short-range and global migration, suggesting that this phenomenon presents a general property of metacommunities and an interesting question for future research. Besides, we also want to mention that, close to the onset of non-zero mean abundances, mean-field theory and numerical solutions are always in very well agreement since there, inter-species interactions are negligible.

## Supplementary Note 8: Variations in growth and dispersal rates of species with global migration

To explore more general metacommunities, where species may differ in their growth, interaction, and migration rates, we generalized Eq. (10) and consider the following dynamics in the metacommunity:

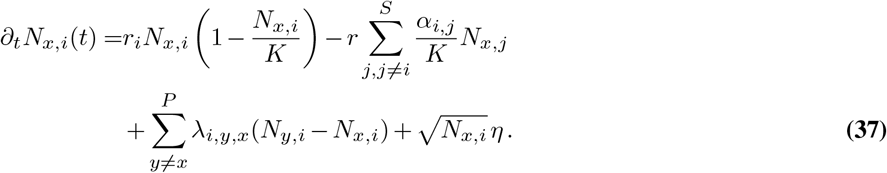

Here, fitness differences between species *i* are implemented by species-specific growth rates *r*_*i*_. Furthermore, the interaction strengths between species, given by *α*_*i,j*_, may differ, and species may have different migration rates *λ*_*i*_. For simplicity, the parameters *r*_*i*_, *λ*_*i*_, and *α*_*i,j*_ are drawn from normal distributions centered around *r, λ*, and *α*, with standard deviations *σ*_*r*_, *σ*_*λ*_, and *σ*_*α*_, respectively (negative migration rates are set to *λ*). With the generalized dynamics Eq. (37), the effective growth factor of a species *i* at *x* is given by 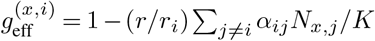. For Fig. 5 in the main text we numerically solved Eq. (37) for short-range migration and relatively small parameter differences across species, in particular 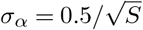 and *σ*_*λ*_ = 0.03 (Fig. 5*A*, main text) and *σ*_*r*_ = 0.03 (Fig. 5*B*, main text). Among other things, we find that at the end of our numerical solutions the patch-averaged effective growth factors 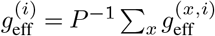 of the surviving species cluster close to the critical threshold value *g*^*c*^.

In the following we will perform a similar analysis for global migration and discuss the effect of different magnitudes of growth parameter variations among the species on the metacommuniy dynamics in more detail. Previous studies on species-rich, well-mixed communities, that ignored demographic fluctuations (14, 16, 17, 50) have shown that when the spread in inter-species interactions *σ*_*α*_ exceeds a certain threshold value 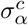 (with all species having the same growth rate, i.e. *σ*_*r*_ = 0), the coexistence state is no longer stable and species will start to go extinct. In this regime, a well-mixed community shows multistability (17), while metacommunities with global migration have been shown to exhibit spatio-temporal chaos (33, 34). Previous studies of metacommunities (33, 34) typically considered a carrying capacity that very large [*K* ∼ 10^9^ in (33)] so that demographic fluctuations can be assumed to play a subordinate role for the dynamics. Furthermore, extinctions are implemented by setting population sizes below one individual to zero (*ad hoc* ‘cutoff’). Following this implementation for large carrying capacities, in Fig. S6*A* we increase the spread *σ*_*α*_ of interaction coefficients and recapitulate a transition from a stable unique state of full coexistence to a state of spatio-temporal chaos even when the species growth rates slightly differ (*σ*_*r*_ = 0.01)^1^. While in Fig. S6*A* we increased the variation in inter-species interactions, since this has been the focus for several previous studies of well-mixed ecosystems (14, 16, 17, 50), we think it will be similarly interesting to investigate a similar transition when increasing variations in other system parameters such as the growth rates as well as migration rates in future studies.

In Fig. S6*B* we return to a regime of smaller carrying capacities (*K* ∼ 1000) and increase the spread *σ*_*α*_ of interaction co-efficients for a metacommunity following Eq. (10), i.e. including demographic fluctuations. When we measure the species’ patch-averaged growth factors 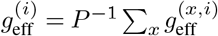, we find that initially these undergo quick relaxation dynamics followed by weak fluctuations. As discussed in the main text (Fig. 2*C*, Fig. 4*A*, Fig. 5*B* in main text), increasing the number of competing species *S* drives the 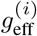 on average closer to the threshold value *g*^*c*^. When we increase the spread *σ*_*α*_ in the species’ interactions coefficients, the spread of the species’ patch-averaged growth factors 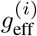 grows [circles in Fig. S6*B*]. For small *σ*_*α*_ [see *σ*_*α*_ = 0.5, 1 in Fig. S6*B*], we observe that only species with patch-averaged effective growth factor 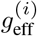 above *g*^*c*^ survive (purple circles), while species with smaller 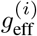 have gone extinct at the end of our numerical solution (gray circles). This is in agreement with our results for short-range migration (see section ‘Variances in growth parameters lead to stochastic extinctions’ in main text and Fig.5). For larger *σ*_*α*_ [see *σ*_*α*_ = 2 in Fig. S6*B*], we find that also species with an patch-averaged effective growth factor 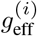 smaller than *g*^*c*^ survive. We hypothesize that the ability of species being able to survive despite of below-threshold effective growth factors (i.e.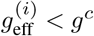), is the multistability present in these regimes of large *σ*_*α*_ [see discussion on multistability above and Refs (14, 16, 17, 50)]. Similar to our discussion in section 3 and section 7, multistability may result in the exclusion among certain communities of species on the same patch and, furthermore, demixing of communities, especially for small migration rates. When different communities demix spatially, the effective growth factor averaged over all patches, *g*^(*i*)^, may no longer be a good indicator for the survival of individual species. This intuition regarding spatial demixing is further supported when we look at the different communities assumed on different patches at different times. To this end we first identified all different community compositions (i.e. the sets of species that have an abundance of at least one individuum) that appear during a numerical solution. Then, we measured how much these communities overlap, i.e. how many species two different communities share. In Fig. S6*C* we plot the distribution of the relative overlap of different identified communities Ω(*x, y, t*), which we define as Ω(*x, y, t*) = |*C*_*x,t*_ ∩ *C*_*y,s*_ |*/*| |*C*_*x,t*_ |, for 100 time samples and all patches. We find that for relatively small variations in the interaction strengths (*σ*_*λ*_ = 0.5, 1 in Fig. S6*C*), the distribution of the relative overlap shows a pronounced peak around closely resembling community, i.e. communities with closely resembling species compositions. In contrast, for larger variations in the interaction strengths (*σ*_*λ*_ = 2 in Fig. S6*C*), there are large contributions from communities with less resemblance (i.e. very different species compositions), suggesting that some communities mutually exclude each other.

We would like to note, that when implementing extinctions by an *ad hoc* ‘cutoff’ instead of explicit demographic noise even for relatively small carrying capacities [e.g. *K* = 1000 as in Fig. S6*B*], we observe that species can in general survive with patch-averaged growth factors 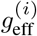 below the threshold value *g*^*c*^ [see Fig. S6*D*]. As discussed in the main text, the need of species to overcome a finite threshold value *g*^*c*^ *>* 0 to survive in our study with explicit demographic noise constitutes an important difference between our work and studies that ignore demographic fluctuations (or implement extinctions by an *ad hoc* ‘cutoff’). While *g*^*c*^ approaches zero for large carrying capacities [see Eq. (35), Eq. (36)], we hypothesize that at smaller population sizes, the need to have a patch-averaged effective growth factor beyond *g*^*c*^ can have important consequences, for instance, on the fixation of an upcoming mutant or a species immigrating from outside the metacommunity.

**Fig. S1.**
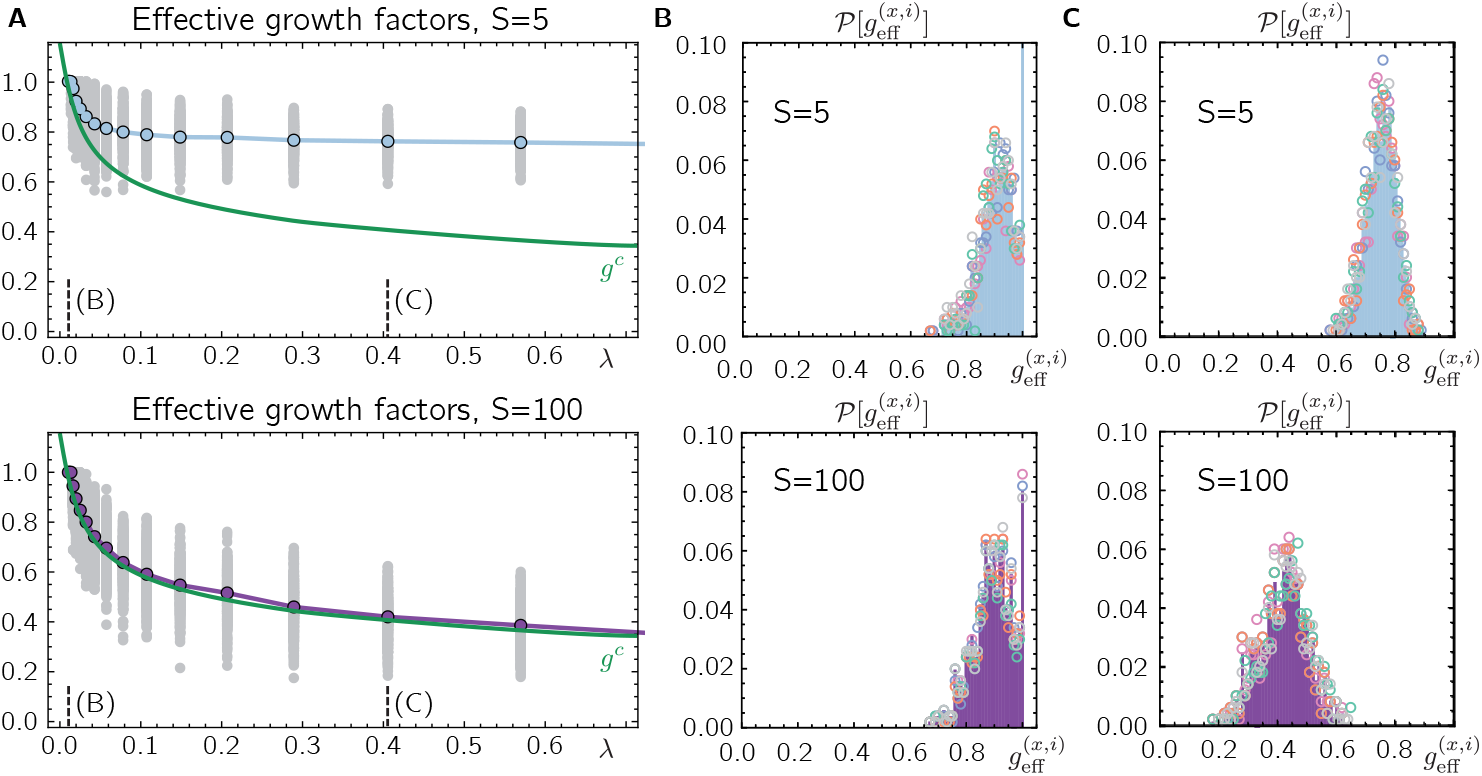
Distribution of effective growth factors. **(A)** Effective growth factors 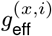 of one representative species for different migration rates and initially coexisting species *S* = 5 (top) and *S* = 100 (bottom). While individual effective growth factors 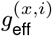 can fall below the critical extinction threshold *g*^*c*^ (green line), the patch-averaged effective growth factors 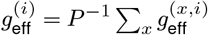 (blue and purple circles for *S* = 5 and *S* = 100, respectively) saturate at the extinction threshold *g*^*c*^ for increasing *S*. **(B), (C)** Distributions of effective growth factors 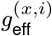 of all species (top: *S* = 5, bottom: *S* = 100) and demes at the final time step of our numerical solution for fixed migration rates [*(B)*: *λ ≈* 0.013 and *(C)*: *λ ≈* 0.4 as indicated by the vertical dashed lines in *(A)*]. Open circles of five different colors denote the distributions of 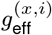 for 5 different species, respectively, which are very similar. Parameters are *r* = 0.3, *K* = 10, *α* = 0.1, *P* = 500.

**Fig. S2.**
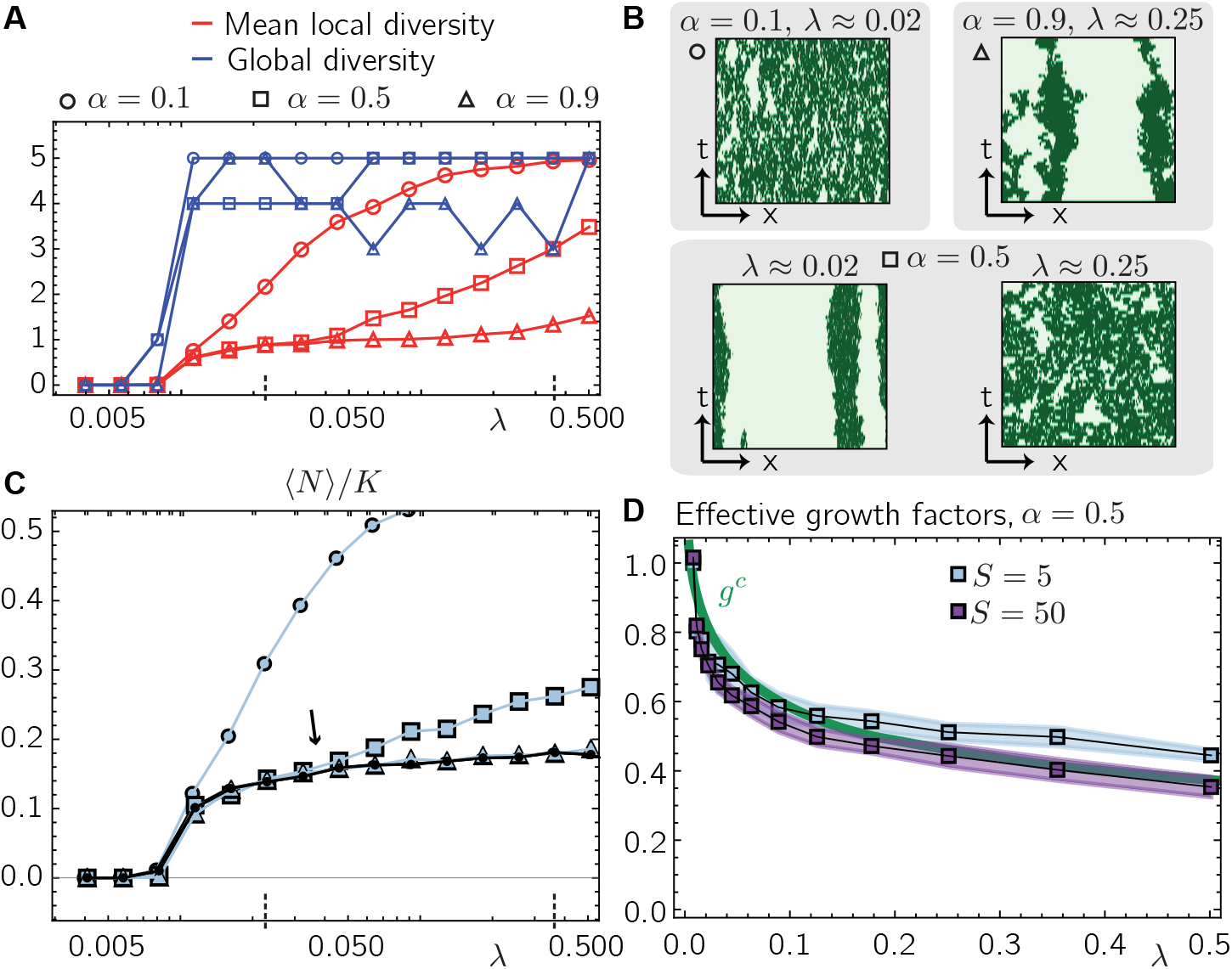
For larger competition strenghts metacommunities with short-range migration show spatial demixing. **(A)** Total number of surviving species in the metacommunity (global diversity, blue symbols) and mean number of surviving species per patch (local diversity, red symbols) for different inter-species competition strengths *α* (circles, squares and triangles denote *α* = 0.1, 0.5 and 0.9, respectively) and migration rates *λ* for *S* = 5. **(B)** Kymographs of individual representative species show that while for weak competitions, species do not demix (top left panel), stronger competition strenghts *α* can lead to spatial demixing already when *α <* 1 even for migration rates *λ* far beyond the threshold value *λ*_*c*_ (i.e. for *λ≈* 0.25 ≫*λ*_*c*_, see top right panel). For intermediate competition strengths (*α* = 0.5, lower panels), we observe a transition between spatial demixing for low migration rates (left) and no demixing for larger migration rates (right). **(C)** Blue shaded symbols denote the mean abundance of all species for different competition strengths *α* (circles, squares and triangles denote *α* = 0.1, 0.5 and 0.9, respectively) as a function of the migration rate *λ*. The black line and circles show the average abundance of a species in the absence of inter-species interactions (i.e. *α* = 0) as a comparison. When the local diversity is close to one [e.g. because of larger *α*, compare (A)], the mean abundance of species follows the abundance of a single species. Here, species spatially demix and do not do not interact on most patches [as indicated in the kymographs in (B)]. When increasing the migration rate *λ*, for intermediate competition strengths (*α* = 0.5, squares) we observe a crossover between a regime of spatial demixing, where the mean abundance follows the one of non-interacting species, and no demixing, where the mean abundance is larger than the one of non-interacting species (the crossover is highlighted by an arrow). **(C)** Mean (solid lines and squares) and across-species deviations (shaded areas) of the patch-averaged effective growth factors 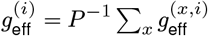 for *S* = 5 (blue) and *S* = 50 (purple) at intermediate competition strengths *α* = 0.5. The demixing suggested in A-C results in stronger across-species variations of the patch-averaged effective growth factors than for weak competitions (see Fig. 2C in the main text). In the regimes where species demix, we see that the measured patch-averaged effective growth factors can fall below *g*^*c*^. This effect of demixing is even more pronounced when the number of species is large (e.g. *S* = 50, purple). The remaining parameters are *r* = 0.3, *K* = 10, *P* = 500.

**Fig. S3.**
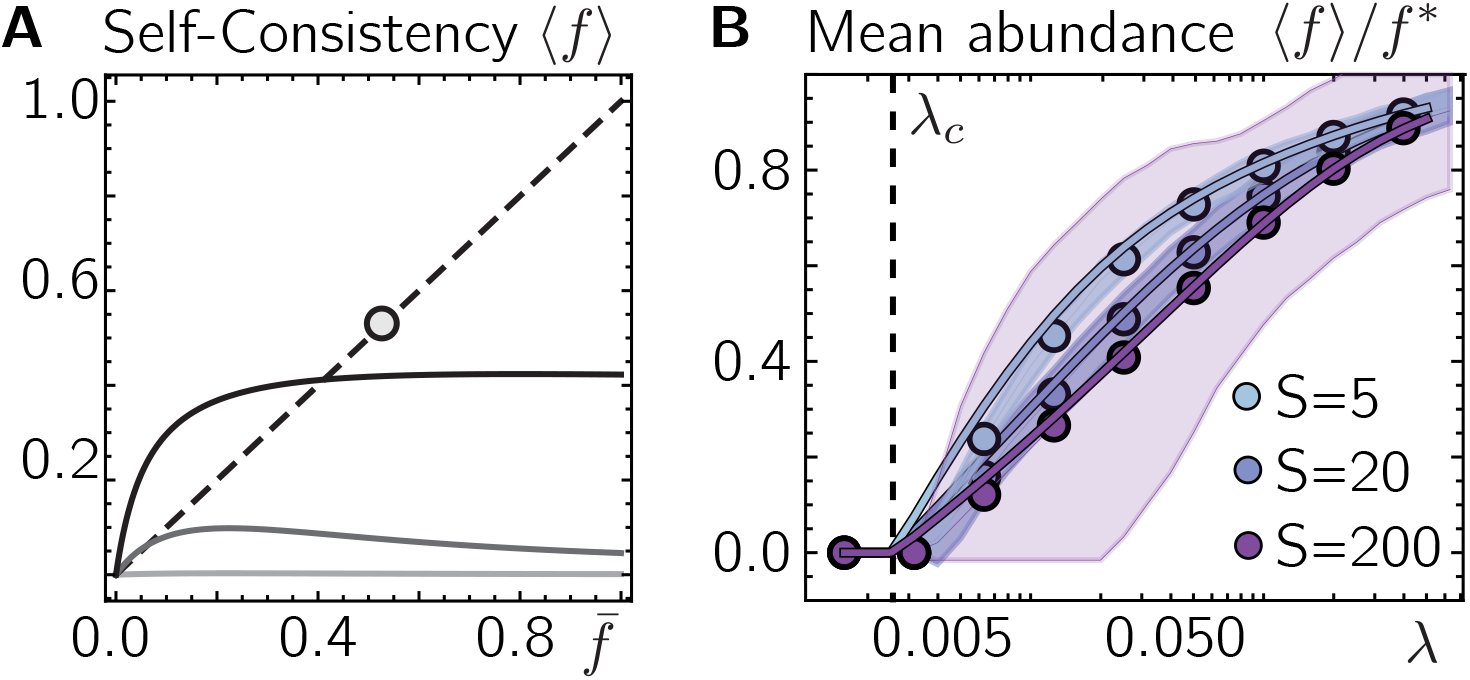
Self-consistent derivation of the mean abundance of species for global migration. **(A)** Above a critical migration rate, *λ*_*c*_, the self-consistency condition 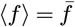, Eq. (24) (dashed line), has a solution with non-zero mean abundance ⟨*f* ⟩ marking an onset of finite population sizes. Shown are the solutions for ⟨*f* ⟩ for migration rates smaller, close above and farther above the critical migration rate *λ*_*c*_ (from bright to dark grey). The circle denotes the deterministic steady stationary solution 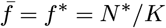. **(B)** The analytic mean field solution for the mean abundance *f)* (solid lines) is in very good agreement with the numerical solution of the explicit multiple species metacommunity dynamics Eq. (18) (circles). The shaded regions denote the standard deviation of the mean abundances ⟨*f*_*i*⟩_ in our numerical solutions across different species, which can be quite large when the number of species *S* is large. The parameters are *r* = 0.3, *K* = 10, *α* = 0.1.

**Fig. S4.**
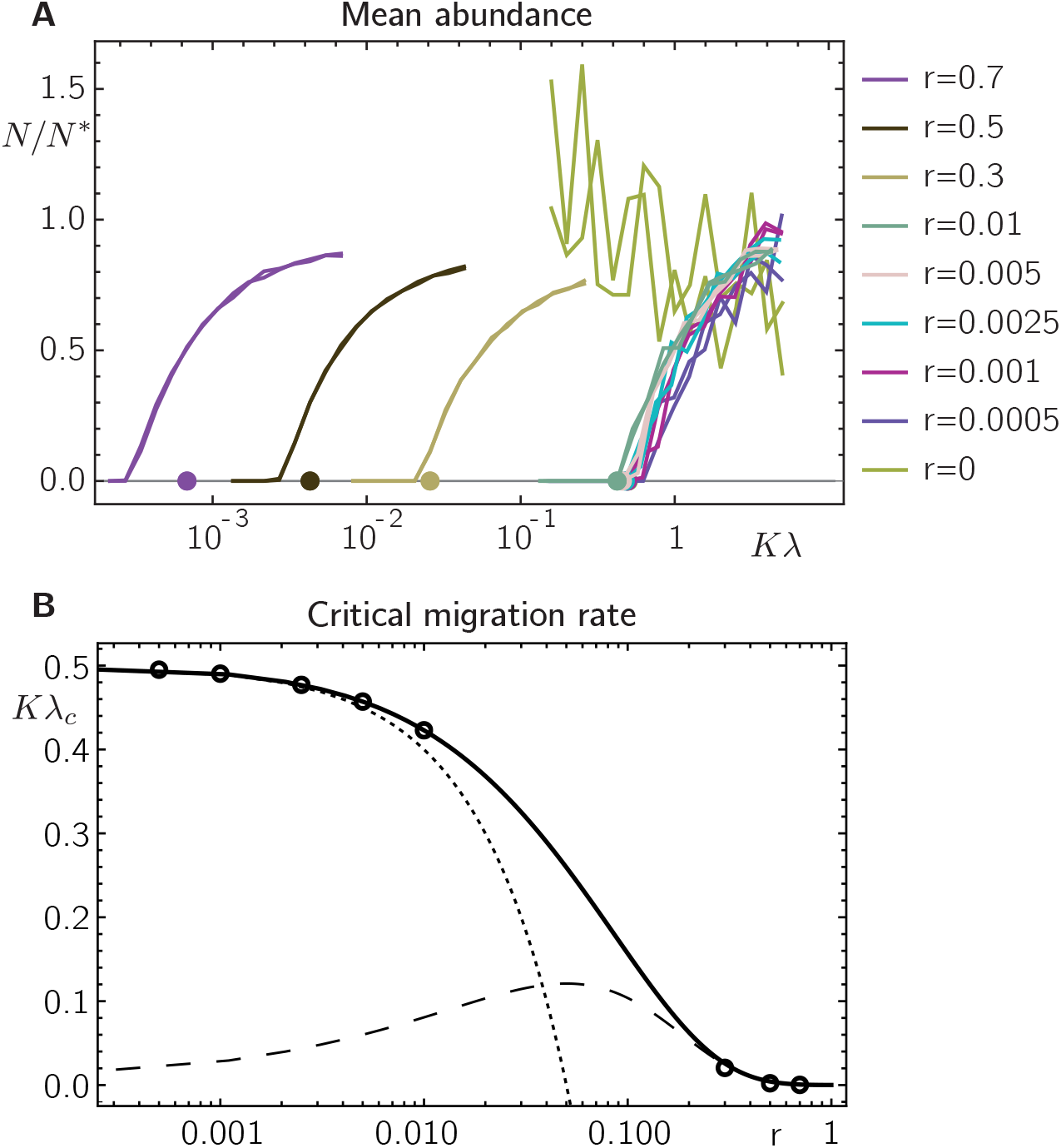
Limiting behaviors of the critical migration rate. **(A)** When decreasing the growth rate *r* in our numerical solutions, the transition from zero to non-zero mean abundances *N/N* ^*^ (solid lines, results for two independent random choices of initial conditions are shown) shifts to larger migration rates *λ* in agreement with our mean-field prediction (circles). As predicted by our mean field theory [see Eq. (27)], the critical migration rate *λ*_*c*_ approaches a finite value for infinitesimally small *r*. Setting *r* = 0, the metacommunity has not yet approached the absorbing state *N*_*x,i*_ = 0 within our simulation time and our numerical results do not suggest a clear dependence of the final mean abundance on the migration rate *λ*. **(B)** The self-consistent mean-field solutions for the critical migration rate *λ*_*c*_(*r*) (solid line) are in very good agreement with our numerical solutions [circles denote the onset migration rates obtained from the numerical solutions in *(A)*]. The dashed and dotted lines denote the limiting behaviors for *Kλ/*(*Kr*) ≪1, Eq. (26), and *Kr/*(*Kλ*) ≪1, Eq. (27), respectively. The remaining parameters are *K* = 10, *α* = 0 and for the numerical solutions: *P* = 2000, Final time: 20000. As initial condition we chose *N*_*x,i*_ = *K* for all patches and species with small random perturbations.

**Fig. S5.**
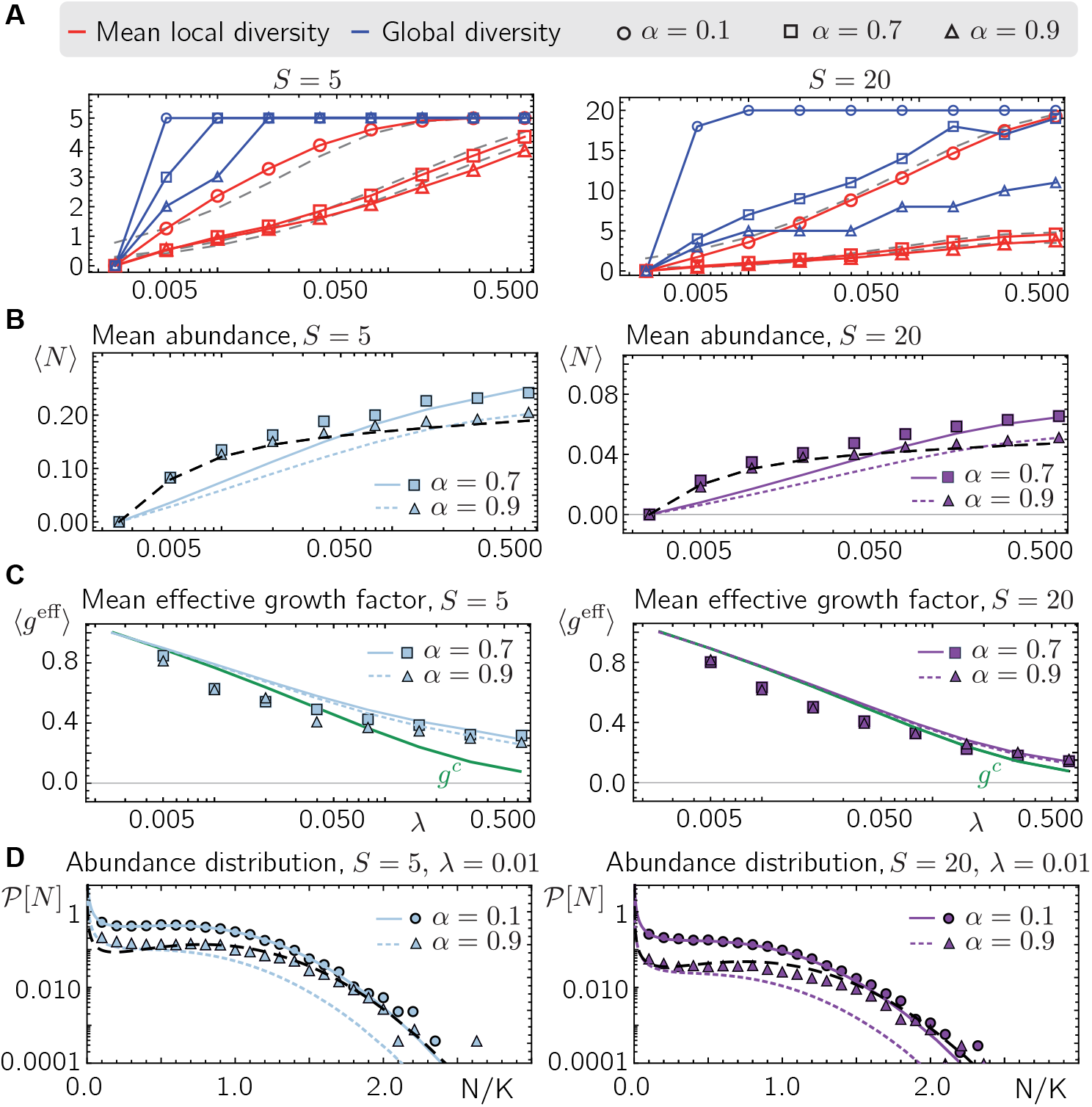
Mean field theory for increasing competition strengths. **(A)** Total number of surviving species in the metacommunity (global diversity, blue symbols and lines) and mean number of surviving species per patch (local diversity, red symbols and solid lines) for different inter-species competition strengths *α* (circles, squares and triangles denote *α* = 0.1, 0.7 and 0.9, respectively) and migration rates *λ* for *S* = 5 (left) and *S* = 20 (right). The dashed gray red lines show the mean local diversity according to the approximation Eq. (30). **(B)** Mean-field predictions (colored lines) and the mean abundance of our numerical solutions (symbols) of all species for different competition strengths *α* (squares and triangles denote *α* = 0.7 and 0.9, respectively) for *S* = 5 (left) and *S* = 20 (right) as a function of the migration rate *λ*. The black dashed line denotes the mean-field prediction for the mean abundance of a species in the absence of inter-species interactions (i.e. *α* = 0) as a comparison. When the local diversity is close to one [e.g. because of larger *α*, compare (A)], the mean abundance of species (i.e. colored lines) follows the abundance of a single species (i.e. black dashed line). When increasing the migration rate *λ*, for intermediate competition strengths (*α* = 0.7, squares) we observe a crossover between a regime where the mean abundance follows the mean-field prediction for non-interacting species, and a regime where the mean abundance is better approximated by the mean-field prediction for species with competition strength *α*. **(C)** Mean-field prediction (left: blue lines for *S* = 5, right: purple lines for *S* = 20) and numerical solutions (respective symbols) for the mean patch-averaged effective growth factors 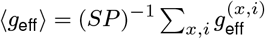 at different competition strengths (squares: *α* = 0.5, triangles: *α* = 0.9). In the regimes where species demix (i.e. the local diversity is typically one), we see that the mean patch-averaged effective growth factor can fall below *g*^*c*^ (green solid lines). **(D)** Mean-field prediction (left: blue lines for *S* = 5, right: purple lines for *S* = 20) and numerical solutions (respective symbols) for the abundance distribution. The black dashed lines denotes *P*[*N*]*/S*, the mean-field prediction for the mean abundance of a single species in a metacommunity of non-interacting species. The remaining parameters are *r* = 0.3, *K* = 10, *P* = 500.

**Fig. S6.**
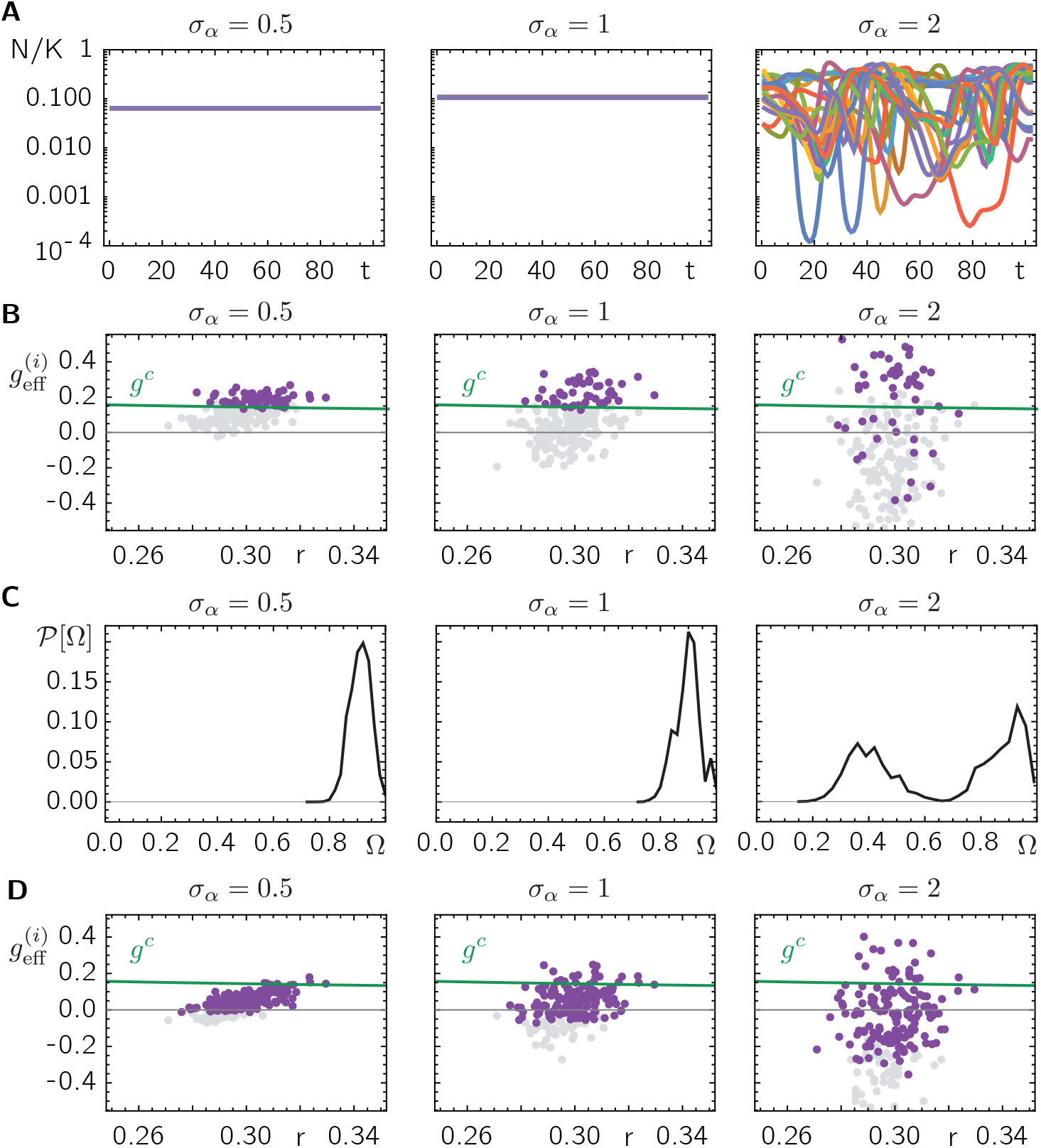
Variations in growth and interaction parameters lead to species extinctions and clustering close to the extinction threshold. **(A)** Abundance of one representative species during hundred time steps for different demes (different colors, every second deme is shown). Increasing the spread *σ*_*α*_ leads to a transition between a stationary state to spatio-temporal chaotic dynamics. For these numerical solutions, demographic fluctuations were ignored and an extinction threshold is implemented by setting population sizes of less than one individual to zero (see section 8). *K* = 10^9^. **(B)** Patch-averaged effective growth factors 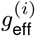 of species at the end of our numerical solution have survived (purple circles) and have gone extinct (gray circles). For *σ*_*α*_ below the transition to multistability indicated in *A*, only species with 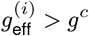 survive, while above the transition also species with 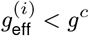 survive, suggesting that in here 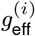 is no longer a good indicator of species survival. *K* = 1000. **(C)** Distribution *P*[Ω] of the overlap Ω (see 8 for definition) of the different communities identified in 100 time samples and all patches. For small *σ*_*α*_, *P*[Ω] has a sharp peak at closely resembling communities (communities with approx 80% overlap); however, for larger *σ*_*α*_ there are additional large contributions in *P*[Ω] from communities with less resemblance (only approx. 40% overlap), suggesting that some communities mutually exclude each other. **(D)** When extinctions are implemented by an *ad hoc* ‘cutoff’ (population sizes of less than one individual are set to zero, see section 8) instead of explicit demographic noise, species can survive even when their patch-averaged growth factor is below the threshold value *g*^*c*^ highlighting an important role of demographic fluctuations. For all solutions in *A*-*D* we used *r* = 0.3, *α* = 0.1, *λ* = 10^−5^, *S* = 200, and *P* = 100.

## Footnotes

1 The critical threshold value for the transition between stable coexistence and multistability in a well-mixed community without differences in the species’ growth and migration rate (i.e. *σ*_*r*_ = *σ*_*λ*_ = 0) is given by 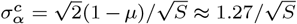 (14, 16, 17, 50), where we accounted for the fact, that the interaction strength in our study does not scale inversely with *S*.

## Bibliography

1. MA Huston. Patterns of species diversity on coral reefs. Annual review of ecology and systematics, 16(1):149–177, 1985.

2. Joseph S Wright. Plant diversity in tropical forests: a review of mechanisms of species coexistence. Oecologia, 130(1):1–14, 2002.

3. Paul B. Rainey and Steven D. Quistad. Toward a dynamical understanding of microbial communities. Philosophical Transactions of the Royal Society B: Biological Sciences, 375 (1798):20190248, 2020.

4. Nadav Kashtan, Sara E Roggensack, Sébastien Rodrigue, Jessie W Thompson, Steven J Biller, Allison Coe, Huiming Ding, Pekka Marttinen, Rex R Malmstrom, Roman Stocker, et al. Single-cell genomics reveals hundreds of coexisting subpopulations in wild prochlorococcus. Science, 344(6182):416–420, 2014.

5. Michael J. Rosen, Michelle Davison, Devaki Bhaya, and Daniel S. Fisher. Fine-scale diversity and extensive recombination in a quasisexual bacterial population occupying a broad niche. Science, 348(6238):1019–1023, 2015.

6. Nandita R Garud, Benjamin H Good, Oskar Hallatschek, and Katherine S Pollard. Evolutionary dynamics of bacteria in the gut microbiome within and across hosts. PLoS biology, 17(1):e3000102, 2019.

7. Robert H MacArthur and Edward O Wilson. An equilibrium theory of insular zoogeography. Evolution, pages 373–387, 1963.

8. Stephen P Hubbell. The Unified Neutral Theory of Biodiversity and Biogeography. Princeton University Press, Princeton, UNITED STATES, 2001. ISBN 9781400837526.

9. David Alonso, Rampal S. Etienne, and Alan J. McKane. The merits of neutral theory. Trends in Ecology and Evolution, 21(8):451–457, 2006.

10. James Rosindell, Stephen Hubbell, and Rampal Etienne. The Unified Neutral Theory of Biodiversity and Biogeography at Age Ten, volume 26. jul 2011.

11. Robert M. May. Will a Large Complex System be Stable? Nature, 238:413–414, 1972.

12. Stefano Allesina and Si Tang. Stability criteria for complex ecosystems. Nature, 483(7388):205–208, 2012.

13. Axel G Rossberg. Food webs and biodiversity: foundations, models, data. John Wiley & Sons, 2013.

14. Guy Bunin. Ecological communities with Lotka-Volterra dynamics. Physical Review E, 95 (4):1–8, 2017.

15. Mikhail Tikhonov and Remi Monasson. Collective Phase in Resource Competition in a Highly Diverse Ecosystem. Physical Review Letters, 118(4):1–5, 2017.

16. Tobias Galla. Dynamically evolved community size and stability of random Lotka-Volterra ecosystems(a). Epl, 123(4):1–13, 2018.

17. Giulio Biroli, Guy Bunin, and Chiara Cammarota. Marginally stable equilibria in critical ecosystems. New Journal of Physics, 20(8), 2018.

18. Mathew A Leibold and Jonathan M Chase. Metacommunity Ecology, Volume 59. Princeton University Press, 2017.

19. Mathew A Leibold, Marcel Holyoak, Nicolas Mouquet, Priyanga Amarasekare, Jonathan M Chase, Martha F Hoopes, Robert D Holt, Jonathan B Shurin, Richard Law, David Tilman, et al. The metacommunity concept: a framework for multi-scale community ecology. Ecology letters, 7(7):601–613, 2004.

20. Diana Fusco, Matti Gralka, Jona Kayser, Alex Anderson, and Oskar Hallatschek. Excess of mutational jackpot events in expanding populations revealed by spatial luria–delbrück experiments. Nature communications, 7:12760, 2016.

21. Oskar Hallatschek, Pascal Hersen, Sharad Ramanathan, and David R Nelson. Genetic drift at expanding frontiers promotes gene segregation. Proceedings of the National Academy of Sciences, 104(50):19926–19930, 2007.

22. Oskar Hallatschek and David R Nelson. Life at the front of an expanding population. Evolution: International Journal of Organic Evolution, 64(1):193–206, 2010.

23. Maxim O Lavrentovich, Mary E Wahl, David R Nelson, and Andrew W Murray. Spatially constrained growth enhances conversional meltdown. Biophysical journal, 110(12):2800–2808, 2016.

24. Jona Kayser, Carl F Schreck, Matti Gralka, Diana Fusco, and Oskar Hallatschek. Collective motion conceals fitness differences in crowded cellular populations. Nature ecology & evolution, 3(1):125–134, 2019.

25. Richard Levins. Some Demographic and Genetic Consequences of Environmental Heterogeneity for Biological Control1. Bulletin of the Entomological Society of America, 15(3):237–240, 09 1969.

26. Ilkka Hanski and Michael Gilpin. Metapopulation dynamics: brief history and conceptual domain. Biological Journal of the Linnean Society, 42(1-2):3–16, 1991.

27. Alan Hastings. Structured models of metapopulation dynamics. Biological Journal of the Linnean Society, 42(1-2):57–71, 1991.

28. JC Allen, WM Schaffer, and D Rosko. Chaos reduces species extinction by amplifying local population noise. Nature, 364(6434):229–232, 1993.

29. Graeme D Ruxton. Low levels of immigration between chaotic populations can reduce system extinctions by inducing asynchronous regular cycles. Proceedings of the Royal Society of London. Series B: Biological Sciences, 256(1346):189–193, 1994.

30. Jordi Bascompte. Extinction thresholds: insights from simple models. In Annales Zoologici Fennici, pages 99–114. JSTOR, 2003.

31. Tobias Reichenbach, Mauro Mobilia, and Erwin Frey. Mobility promotes and jeopardizes biodiversity in rock–paper–scissors games. Nature, 448(7157):1046–1049, 2007.

32. Yossi Ben-Zion and Nadav M. Shnerb. Coherence, conservation and patch-occupancy analysis. Oikos, 121(6):985–997, 2012.

33. Michael T Pearce, Atish Agarwala, and Daniel S Fisher. Stabilization of extensive fine-scale diversity by ecologically driven spatiotemporal chaos. Proceedings of the National Academy of Sciences, 117(25):14572–14583, 2020.

34. Felix Roy, Matthieu Barbier, Giulio Biroli, and Guy Bunin. Complex interactions can create persistent fluctuations in high-diversity ecosystems. PLoS computational biology, 16(5): e1007827, 2020.

35. Peter Chesson. Macarthur’s consumer-resource model. Theoretical Population Biology, 37 (1):26–38, 1990.

36. Peter Chesson. Mechanisms of Maintanance of species diversity. Annual Review of Ecology and Systematics, 31:343–66, 2000.

37. Michaela M Salcher. Same same but different: ecological niche partitioning of planktonic freshwater prokaryotes. Journal of Limnology, 73(1s):74–87, 2014.

38. Deborah L Finke and William E Snyder. Niche partitioning increases resource exploitation by diverse communities. Science, 321(5895):1488–1490, 2008.

39. EM Chávez-Solís, C Solís, N Simões, and M Mascaró. Distribution patterns, carbon sources and niche partitioning in cave shrimps (atyidae: Typhlatya). Scientific reports, 10(1):1–16, 2020.

40. Richard Baran, Eoin L Brodie, Jazmine Mayberry-Lewis, Eric Hummel, Ulisses Nunes Da Rocha, Romy Chakraborty, Benjamin P Bowen, Ulas Karaoz, Hinsby Cadillo-Quiroz, Ferran Garcia-Pichel, et al. Exometabolite niche partitioning among sympatric soil bacteria. Nature communications, 6:8289, 2015.

41. Aaron Goodman and Lauren Esposito. Niche partitioning in congeneric scorpions. Invertebrate Biology, 139(1):e12280, 2020.

42. Haldre S Rogers, Noelle G Beckman, Florian Hartig, Jeremy S Johnson, Gesine Pufal, Katriona Shea, Damaris Zurell, James M Bullock, Robert Stephen Cantrell, Bette Loiselle, Liba Pejchar, Onja H Razafindratsima, Manette E Sandor, Eugene W Schupp, W Christopher Strickland, and Jenny Zambrano. The total dispersal kernel: a review and future directions. AoB PLANTS, 11(5), 09 2019.

43. Robert MacArthur and Richard Levins. The limiting similarity, convergence, and divergence of coexisting species. The american naturalist, 101(921):377–385, 1967.

44. Bart Haegeman and Rampal S. Etienne. Self-consistent approach for neutral community models with speciation. Physical Review E - Statistical, Nonlinear, and Soft Matter Physics, 81(3):1–13, 2010.

45. David A. Kessler and Nadav M. Shnerb. Generalized model of island biodiversity. Physical Review E - Statistical, Nonlinear, and Soft Matter Physics, 91(4):1–11, 2015.

46. Ricky Der, Charles L Epstein, and Joshua B Plotkin. Generalized population models and the nature of genetic drift. Theoretical population biology, 80(2):80–99, 2011.

47. Maxim O Lavrentovich, Kirill S Korolev, and David R Nelson. Radial domany-kinzel models with mutation and selection. Physical Review E, 87(1):012103, 2013.

48. Haye Hinrichsen. Non-equilibrium critical phenomena and phase transitions into absorbing states. Advances in Physics, 49(7):815–958, 2000.

49. Géza Ódor. Universality classes in nonequilibrium lattice systems. Reviews of modern physics, 76(3):663, 2004.

50. Felix Roy, Giulio Biroli, Guy Bunin, and Chiara Cammarota. Numerical implementation of dynamical mean field theory for disordered systems: Application to the lotka–volterra model of ecosystems. Journal of Physics A: Mathematical and Theoretical, 52(48):484001, 2019.

51. Mehran Kardar. Statistical physics of fields. Cambridge University Press, 2007.

52. Greg Huber, Mogens H Jensen, and Kim Sneppen. Distributions of self-interactions and voids in (1+ 1)-dimensional directed percolation. Physical Review E, 52(3):R2133, 1995.

53. Ronald Dickman and Marcelo Martins de Oliveira. Quasi-stationary simulation of the contact process. Physica A: Statistical Mechanics and its Applications, 357(1):134–141, 2005.

54. Ran Nathan. Long-distance dispersal of plants. Science, 313(5788):786–788, 2006.

55. Joshua B Plotkin, Matthew D Potts, W Yu Douglas, Sarayudh Bunyavejchewin, Richard Condit, Robin Foster, Stephen Hubbell, James LaFrankie, N Manokaran, Lee Hua Seng, et al. Predicting species diversity in tropical forests. Proceedings of the National Academy of Sciences, 97(20):10850–10854, 2000.

56. Igor Volkov, Jayanth R Banavar, Stephen P Hubbell, and Amos Maritan. Patterns of relative species abundance in rainforests and coral reefs. Nature, 450(7166):45–49, 2007.

57. Igor Volkov, Jayanth R Banavar, Stephen P Hubbell, and Amos Maritan. Neutral theory and relative species abundance in ecology. Nature, 424(6952):1035–1037, 2003.

58. Sz Horvát, A Derzsi, Z Néda, and A Balog. A spatially explicit model for tropical tree diversity patterns. Journal of theoretical biology, 265(4):517–523, 2010.

59. Alan McKane, David Alonso, and Ricard V. Solé. Mean-field stochastic theory for speciesrich assembled communities. Physical Review E - Statistical Physics, Plasmas, Fluids, and Related Interdisciplinary Topics, 62(6 B):8466–8484, 2000.

60. Ricard V. Solé, David Alonso, and Alan McKane. Self-organized instability in complex ecosystems. Philosophical Transactions of the Royal Society B: Biological Sciences, 357 (1421):667–681, 2002.

61. Bart Haegeman and Michel Loreau. A mathematical synthesis of niche and neutral theories in community ecology. Journal of Theoretical Biology, 269(1):150–165, 2011.

62. Evelyn C Pielou et al. An introduction to mathematical ecology. An introduction to mathematical ecology., 1969.

63. Ganapati P Patil, Evelyn Chris Pielou, William E Waters, WR Waters, and William Alexander Waters. Statistical ecology: spatial patterns and statistical distributions, volume 1. Penn State University Press, 1971.

64. Ricard V. Solé and Susanna C. Manrubia. Extinction and self-organized criticality in a model of large-scale evolution. Physical Review E - Statistical Physics, Plasmas, Fluids, and Related Interdisciplinary Topics, 54(1):R42–R45, 1996.

65. Rampal S. Etienne and David Alonso. Neutral community theory: How stochasticity and dispersal-limitation can explain species coexistence. Journal of Statistical Physics, 128(1-2):485–510, 2007.

66. Graham Bell. The distribution of abundance in neutral communities. The American Naturalist, 155(5):606–617, 2000.

67. Rick Durrett and Simon Levin. Spatial models for species-area curves. Journal of Theoretical Biology, 179(2):119–127, 1996.

68. Evan P Economo and Timothy H Keitt. Species diversity in neutral metacommunities: a network approach. Ecology letters, 11(1):52–62, 2008.

69. Patrick B Warren. Biodiversity on island chains: neutral model simulations. Physical Review E, 82(5):051922, 2010.

70. Guido Van Rossum and Fred L Drake Jr. Python reference manual. Centrum voor Wiskunde en Informatica Amsterdam, 1995.

71. Wolfram Research, Inc. Mathematica, Version 12.3.1. Champaign, IL, 2021.

72. Crispin W Gardiner et al. Handbook of stochastic methods, volume 3. springer Berlin, 1985.

73. Joachim Mathiesen, Namiko Mitarai, Kim Sneppen, and Ala Trusina. Ecosystems with mutually exclusive interactions self-organize to a state of high diversity. Physical review letters, 107(18):188101, 2011.

74. James Franklin Crow, Motoo Kimura, et al. An introduction to population genetics theory. An introduction to population genetics theory., 1970.

75. Yoh Iwasa. Free fitness that always increases in evolution. Journal of Theoretical Biology, 135(3):265–281, 1988.

76. N. H. Barton and H. P. De Vladar. Statistical mechanics and the evolution of polygenic quantitative traits. Genetics, 181(3):997–1011, 2009.

77. Ronald Dickman and Alex Yu Tretyakov. Hyperscaling in the domany-kinzel cellular automaton. Physical Review E, 52(3):3218, 1995.

78. Miguel A Munoz, G Grinstein, and Yuhai Tu. Survival probability and field theory in systems with absorbing states. Physical Review E, 56(5):5101, 1997.

79. R. A. Fisher, A. Steven Corbet, and C. B. Williams. The Relation Between the Number of Species and the Number of Individuals in a Random Sample of an Animal Population. The Journal of Animal Ecology, 12(1):42, 1943.

80. Steinar Engen and Russell Lande. Population dynamic models generating species abundance distributions of the gamma type. Journal of Theoretical Biology, 178(3):325–331, 1996.

81. Rampal S. Etienne and David Alonso. A dispersal-limited sampling theory for species and alleles. Ecology Letters, 8(11):1147–1156, 2005.

